# Retinoblastoma-related (RBR) has both canonical and non-canonical regulatory functions during thermo-morphogenic responses in Arabidopsis seedlings

**DOI:** 10.1101/2024.05.28.596227

**Authors:** Rasik Shiekh Bin Hamid, Fruzsina Nagy, Nikolett Kaszler, Ildikó Domonkos, Magdolna Gombos, Eszter Molnár, Aladár Pettkó-Szandtner, László Bögre, Attila Fehér, Zoltán Magyar

## Abstract

Warm temperatures accelerate plant growth, but the underlying molecular mechanism is not fully understood. Here, we show that increasing the temperature from 22°C to 28°C rapidly activates proliferation in the apical shoot and root meristems of wild-type Arabidopsis seedlings. We found that one of the central regulators of cell proliferation, the cell cycle inhibitor RETINOBLASTOMA-RELATED (RBR), is suppressed by warm temperatures. RBR became hyper-phosphorylated at a conserved CYCLIN-DEPENDENT KINASE (CDK) site in young seedlings growing at 28°C, in parallel with the stimulation of the expressions of the regulatory CYCLIN D/A subunits of CDK(s). Interestingly, while under warm temperatures ectopic RBR slowed down the acceleration of cell proliferation, it triggered elongation growth of post-mitotic cells in the hypocotyl. In agreement, the central regulatory genes of thermomorphogenic response, including *PIF4* and *PIF7*, as well as their downstream auxin biosynthetic *YUCCA* genes (*YUC1-2* and *YUC8-9*) were all up-regulated in the ectopic RBR expressing line but down-regulated in a mutant line with reduced RBR level. We suggest that RBR has both canonical and non-canonical functions under warm temperatures to control proliferative and elongation growth, respectively.

## Introduction

Climate change is a major concern of current and future crop production efficiency and food security (Ray *et al*., 2019; Anderson et al., 2020; Ortiz-Bobea *et al*., 2021; Mirón *et al*., 2023). Besides the expected increase in the frequency of extreme weather conditions (Arnell *et al*., 2019; Mirón *et al*., 2023), the steady rise in the average global temperature is also a challenge.

The non-stressful but significant increase in average temperatures, not only accelerates the metabolism and growth of plants, but also activates adaptation processes known as thermomorphogenesis (Casal and Balasubramanian, 2019). Typical features of thermomorphogenesis include the elongation of hypocotyl, stem, petiole, as well as the hyponasty of leaves with decreased surface area (Quint *et al*., 2016; Casal and Balasubramanian, 2019; Lippmann *et al*., 2019; Wang and Zhu, 2022), which altogether contribute to enhanced evaporative leaf cooling (Crawford *et al*., 2012; Bridge *et al*., 2013; Park *et al*., 2019). The elongation of the primary root, along with alterations in the overall root architecture, is now recognized as a component of the thermomorphogenic response, albeit still with limited understanding in this area (Fonseca de Lima *et al*., 2021).

It has become apparent that in the aboveground organs, ambient light and temperature signalling share common pathways (Casal and Qüesta, 2018; Romero-Montepaone *et al*., 2021; Qi *et al*., 2022; Wang and Zhu, 2022; Legris, 2023; Wu *et al*., 2023). The photoreceptor Phytochrome B (PhyB) was found to act also as a thermosensor in plants (Jung *et al*., 2016). It has been demonstrated that PhyB can discriminate between light and temperature cues due to conformation changes and phase separation, respectively (Chen *et al*., 2022). Parallel light- and thermosensing by photoreceptors is integrated by the PHYTOCHROME INTERACTING FACTOR 4 (PIF4) transcription factor (Koini *et al*., 2009; Stavang *et al*., 2009; Hayes et al., 2021; Bian *et al*., 2022; Jung et al., 2023). PIF4 is regulated both transcriptionally and post-translationally in response to light and thermal signals, which fine-tune its level and activity (Ma *et al*., 2016; Martínez *et al*., 2018; Han *et al*., 2019; Lee *et al*., 2020; Qiu, 2020; Qiu *et al*., 2021; Saitoh *et al*., 2021; Xu and Zhu, 2021; Zhao and Bao, 2021; Verma *et al*., 2023, Preprint). During ambient temperature-dependent hypocotyl elongation, auxin synthesis and signalling are among the central targets of PIF4 (Koini *et al*., 2009; Franklin *et al*., 2011; Sun *et al*., 2012; Bianchimano *et al*., 2023). PIF4 directly up-regulates, among others, the genes coding YUCCA8 (YUC8), INDOLE-3-ACETIC ACID INDUCIBLE 19 (IAA19), and INDOLE-3-ACETIC ACID INDUCIBLE 29 (IAA29). YUC8 is a rate-limiting auxin biosynthesis enzyme that has central role in temperature-dependent hypocotyl elongation (Franklin *et al*., 2011; Sun *et al*., 2012). In Arabidopsis, besides those dependent on phyB, temperature sensing mechanisms involving the thermosensory PHYTOCHROME INTERACTING FACTOR 7 (PIF7) and EARLY FLOWERING3 (ELF3) TFs have also been uncovered (Box *et al*., 2015; Raschke *et al*., 2015; Press *et al*., 2016; Chung *et al*., 2020; Fiorucci *et al*., 2020; Burko *et al*., 2022). Both PIF7 and ELF3 have been reported to modulate the function of PIF4 at elevated temperatures (Box *et al*., 2015; Raschke *et al*., 2015; Chung *et al*., 2020; Fiorucci *et al*., 2020; Burko *et al*., 2022). In addition, various epigenetic mechanisms add to the complexity of thermomorphogenesis regulation (Perrella *et al*., 2022).

In comparison to the hypocotyl, which has been thoroughly characterized in this respect, the thermomorphogenic response of the root is less explored (Fonseca de Lima *et al*., 2021). Under laboratory conditions, moderate warmth of 26-29°C is enough to stimulate the growth of the primary root of Arabidopsis, a phenomenon that is mainly controlled by auxin (Fonseca de Lima *et al*., 2021; Ai *et al*., 2023; Bianchimano *et al*., 2023), although in long-term responses the primary role of brassinosteroids has been implicated (Martins *et al*., 2017). Results from Gaillochet et al. (Gaillochet *et al*., 2020) and Borniego et al. (Borniego *et al*., 2022) illustrate that phyB, ELF3, and PIF7 are not responsible for root thermosensing, though their action in the shoot might indirectly influence root growth. While the thermo-regulated elongation of the Arabidopsis hypocotyl primarily depends on auxin- and brassinosteroid-dependent increases in cell size, it is less dependent on cell division (Bellstaedt *et al*., 2020). However, recent study has reported that in the thermosensory response of the roots, an auxin-mediated increase in cell division activity in the meristem is primary and predominant (Ai *et al*., 2023). One may ask if warmer temperatures affect the cell division rate in the meristematic region of the root, do they have the same effect in that of the shoot? Interestingly, there is no research on the thermoresponsiveness of the shoot meristem at the seedling stage.

Thus, in the root, we do not know how the thermal regulation of auxin signalling is linked to the observed cell division response, while in the shoot, the contribution of accelerated cell divisions to the thermomorphogenic response is fully disregarded. If warming has a specific and positive effect on meristem activity, it likely concerns the increased rate of cells entering the cell cycle. Cell cycle entry is controlled by evolutionarily conserved molecular mechanisms, including the RETINOBLASTOMA (RB) protein, which negatively regulates cell cycle progression through its interaction with the E2-promoter binding transcription factor (E2F, Henley and Dick, 2012). Arabidopsis has a single gene coding for the RETINOBLASTOMA-RELATED (RBR) protein, which is implicated in the integration of environmental signals with cell proliferation and differentiation (Harashima and Sugimoto, 2016). The RBR protein is essential for meristem integrity and function, as well as for lateral organ formation (Borghi *et al*., 2010; Desvoyes & Gutierrez, 2020). The widely-accepted view is that RBR is regulated through phosphorylation by CYCLIN-DEPENDENT KINASE (CDK)-CYCLIN D (CYCD) complexes in response to mitogenic signals. This releases E2F-DIMERIZATION PARTNER (DP) heterodimeric transcription factors from RBR-dependent inhibition facilitating cell cycle entry (Sablowski and Gutierrez, 2022).

However, RBR is not only an inhibitor of the cell cycle, but also required for establishing and maintaining quiescence and the correct differentiation of cells leaving the meristem (Wyrzykowska *et al*., 2006; Borghi *et al*., 2010; Perilli *et al*., 2013; Harashima and Sugimoto, 2016; Gombos *et al*., 2023). Interestingly, RBR controls quiescence in complex with canonical E2Fs in both mitotically less active stem cells and post-mitotic differentiated cells by co-repressing cell cycle genes (Gombos et al., 2023). Based on this, the RBR protein is a suitable subject to study whether increased ambient temperature affects the balance between cell division and differentiation. Here we report the thermomorphogenic response of Arabidopsis plants with distinct RBR protein levels highlighting the potential role of this multitasking protein not only in the control of temperature-dependent meristematic cell divisions but also in the elongation of differentiating cells. Interestingly, we have detected different RBR functions in the proliferative meristematic shoot and root apices, as well as in the hypocotyls. Its canonical cell cycle inhibitory function was repressed by warm temperatures in the meristems, while it has a non-canonical activator role on regulatory genes of thermo-morphogenic cell elongation in the hypocotyl.

## Results

### Cell proliferation is activated by warm temperatures in the meristems of young Arabidopsis seedlings

Rising ambient temperatures accelerate plant growth and development, but the underlying mechanisms are not completely elucidated. Plant growth heavily relies on apical meristems, which are the main sites of cell proliferation in plants. Therefore, it is astonishing that the impact of elevated temperatures on cell proliferation in shoot and root apices remains largely unexplored.

To understand the early events triggered by warm temperatures in Arabidopsis plants, young seedlings grown under long-day conditions were examined after being transferred from 22°C to 28°C four days after germination (4DAG). Two days later, we investigated the effect of the temperature increase on the activity of apical meristems. Examining the leaf primordia in the shoot apex using a scanning electron microscope (SEM), the size of the primordia was enlarged at 28°C compared to those developed at 22°C (Fig. 1A). This showed that warm temperatures quickly accelerated primordium development. Measurements of epidermal cell size in the 3^rd^ leaf primordium of seedlings grown at either 22°C or 28°C revealed no apparent differences, falling within the range of 20-60 µm characteristic of meristemoid and protodermal cells (Fig. 1B, Dong et al., 2009). These data support the idea that warm temperatures accelerate primordium growth by activating cell proliferation rather than cell elongation. The faster development of seedlings was even more evident five days after being transferred to the warmer temperature. At this time, the seedlings that were developed at 28°C had four visible-sized leaves, while those grown at 22°C only had two (Fig. 1C). In addition, the first leaf pair (L1-2) was significantly larger at 28°C than at 22°C (Fig. 1D,E). To investigate the reason for their enlargement, we measured the area of palisade leaf cells and found that they were nearly 1.5 times larger at 28°C than at 22°C (Fig. 1F). By calculating the number of constituent cells in these leaves, we discovered that it was doubled at 28°C compared to 22°C (Fig. 1G). These data indicate that both cell proliferation and cell enlargement were activated by warm temperatures during early leaf development, resulting in an enlarged leaf area.

**Figure 1.**
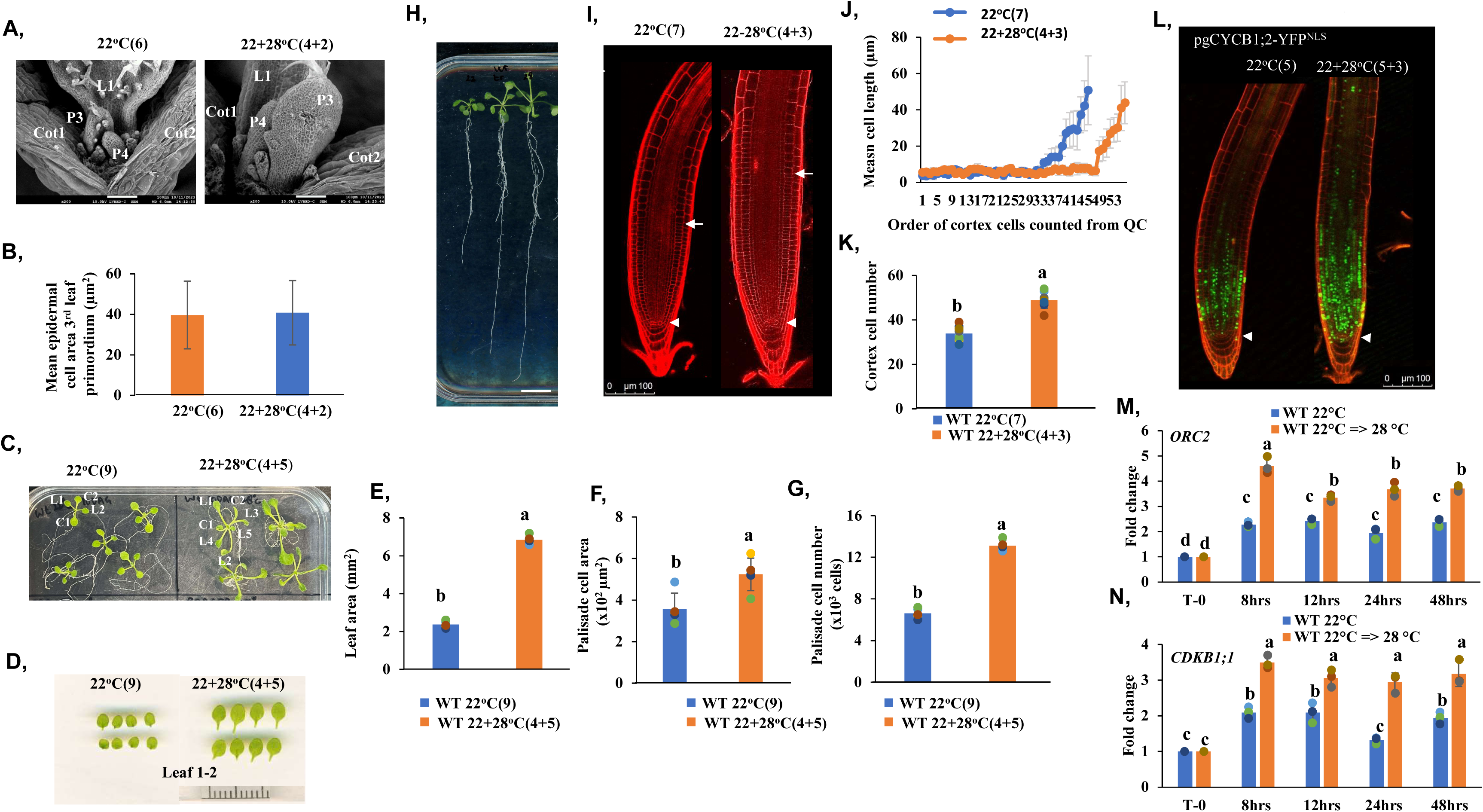
Warm temperatures increase proliferation activity in both the shoot and root meristems of Arabidopsis seedlings. Wild-type seedlings were grown continuously at 22°C (control) or transferred from 22°C to 28°C (22+28°C) for the number of days indicated in the parentheses. **A**) Leaf primordia of seedlings that were grown at a continuous 22°C for six days after germination (DAG) 22°C(6) or two days after transfer of four-day-old seedlings from 22°C to 28°C (22+28°C(4+2)) are pictured under a scanning electron microscope. Positions and orders of cotyledons (Cot), leaves (L), and primordia (P) are indicated by numbers. Scale bar is 100 µm. **B**) The epidermal cell size of the 3^rd^ leaf primordium was determined by using Image J software. **C**) The developmental order of cotyledons (C) and leaves (L) is marked with numbers on seedlings grown at 22°C(9) and 22+28°C(4+5). **D**) Representative pictures of the first leaf pairs of seedlings grown at 22°C(9) or at 22+28°C(4+5). **E-G**) Leaf area (**E**), palisade cell area (**F**), and palisade cell number (**G**) were determined for the first leaf of seedlings grown at the indicated temperature regimes. **H**) The seedlings from left to right were grown at 22°C(8), 22+28°C(4+4), or 28°C(8), respectively. Scale bar is 1 cm. **I**) Representative pictures of propidium iodide-stained roots of seedlings grown at a continuous 22°C(7) or 22+28°C(4+3) obtained by confocal laser scanning microscopy. Arrowheads indicate the quiescent centers, and arrows point to the first elongated cells in the cortical cell file, reflecting meristem size. Scale bar is: 100 µm. **J**) Spatial distribution of cell length across root meristems in seedlings grown at continuous 22°C(7) or 22+28°C(4+3). Mean cell length was calculated at each position along the cortical cell file. The positions were defined by counts of cortical cells from the quiescent centre. Values are averages from the data obtained by the analysis of eight roots from different plants (±SD). **K**) Cortex cell number in the same roots as in J. **L**) The root tips of transgenic seedlings expressing CYCB1;2-YFP^NLS^, a G2 and M-phase marker, under the control of its own promoter. The seedlings were grown at 22°C(6) or 22+28°C(4+2), respectively, before being stained with propidium iodide and imaged using a confocal laser scanning microscope. Arrowheads indicate the quiescent centres. Scale bar is: 100µm. **M-N**) The expression of the S- and the G2-M-phase regulatory genes, *ORC2* (**M**) and *CDKB1;1* (**N**), is elevated in seedlings transferred at four DAG from 22°C to 28°C compared to seedlings grown continuously at 22°C. Expressions were analyzed at 8,12, 24, and 48 hours (hrs) after the time of transfer in seedlings at both 22°C and 28°C, respectively. Values represent fold change normalized to the value of the relevant transcript of the seedlings at T0 (four DAG), which was set arbitrarily at 1. Data are means +/- SD of three biological replicates. Different letters indicate significant differences (P < 0.05) based on one-way ANOVA analyses with Tukey’s HSD post-hoc test (**B, E-K, M, N**).

It is well-known that warm temperatures stimulate root growth resulting in much longer roots at 28°C than at 22°C (Quint et al., 2005; Quint et al., 2009; Hanzawa et al., 2013; Wang et al., 2016; Feraru et al., 2019; Galliochet et al., 2020, Ai et al., 2023). When the seedlings were continuously grown at 28°C, their roots were the longest (Fig. 1H, and Supplemental Fig. S1A). Warm temperatures also resulted in significantly longer roots when the seedlings were transferred from 22°C to 28°C at 4DAG for a period of five more days (Fig. 1H and Supplemental Fig. S1A). To investigate why roots at 28°C grow longer than at 22°C, the root tips of seven-day-old seedlings either grown at 22°C continuously or transferred from 22°C to 28°C at 4DAG for 3 more days were stained with propidium iodide (PI). The samples were then analysed under confocal microscopy (Fig. 1I). The length of cortex cells was determined and plotted based on their distance from the quiescent centre. The results showed that warm temperatures increased the size of the cells (Supplemental Fig. S1B) and elevated the number of small, non-elongated cells in the cortical cell file, indicating that the size of the root meristem enlarged at the higher temperature (Fig. 1I,J,K). To further support this assumption, we grew Arabidopsis seedlings expressing the mitotic marker CYCB1;2 under the control of its own promoter and fused with YFP containing a nuclear localization signal (Iwata et al., 2013). We then compared the fluorescence signal in the roots grown either at 22°C or after being transferred from 22°C to 28°C for 1 or 2 days (Supplemental Fig. S1C and Fig. 1L). We detected an expanded CYCB1;2-YFP signal in the roots due to their growth at 28°C, confirming that more root cells were in mitosis in the root meristem after the temperature was switched from 22°C to 28°C. Finally, we analyzed the expression of two cell cycle regulatory genes: the S-phase specific *ORIGIN RECOGNITION COMPLEX 2* (*ORC2*), and the G2-M-phase specific *CYCLIN-DEPENDENT KINASE B1;1* (*CDKB1;1*) in seedlings 8, 24 and 48 hours after transferring them from 22°C to 28°C (Fig. 1M-N). We compared these expression data with those obtained from seedlings continuously grown at 22°C. Both cell cycle regulatory genes were up-regulated in the first sample, only 8 hours after the seedlings were transferred from 22°C to 28°C. The expression of the genes remained at a higher level throughout the 28°C treatment compared to the continuous growth at 22°C, further supporting that warm temperatures activate cell proliferation.

All together, these data show that raising the ambient temperature from 22°C to 28°C rapidly activates cell proliferation in both the shoot and root meristems.

### The function of the RETINOBLASTOMA-RELATED (RBR) cell cycle inhibitor is suppressed by warm temperatures, in correlation with its phosphorylation status

It is well established that cell proliferation in plants is regulated by the RETINOBLASTOMA-RELATED protein coded by a single gene in Arabidopsis (Desvoyes & Gutierrez, 2020). We assumed that warm temperatures might regulate cell proliferation by controlling the function of RBR. We were curious about whether RBR is indeed involved in this regulation and if yes, how. Previously, we have generated a transgenic Arabidopsis line expressing RBR under the control of its own promoter (Magyar et al., 2012). The protein was fused to a GFP-tag providing an opportunity to follow RBR level in seedlings growing at different temperatures. Confirming previous findings, RBR is expressed at a ubiquitously high level in the root tip of young seedlings at 22°C (Supplemental Fig. S1D). When seedlings were grown at a continuous temperature of 22°C or 28°C for 5 days after germination or transferred to 28°C 4DAG for 8 or 24 hours, respectively, the GFP signal was present both in the root tip and the hypocotyl (Supplemental Fig. S1D_F). Beside the root meristem, strong RBR-GFP signal was detected in non-dividing post-mitotic cells in the transient zone of the root as well as in the hypocotyl epidermis both at 22°C and at 28° C (Supplemental Fig. S1D_F). These data indicated that the RBR protein accumulates in the root and hypocotyl at both 22°C and 28°C. Therefore, it may be involved in regulating both cell proliferation and cell enlargement in response to warm temperatures.

Previously, the RBR-GFP fusion under the control of its own promoter was reported to be functional resulting in a slightly smaller root meristem when expressed in wild-type seedlings (Magyar et al., 2012). Using a western blot assay, we confirmed an increased RBR level in this transgenic line compared to the WT (Supplemental Fig. S1G). In contrast, slightly less RBR protein was detected in the *rbr1-2* mutant seedlings than in the WT confirming previous observations (Chen et al, 2011). The modest reduction in RBR levels resulted in an over-proliferation of meristemoid-like cells in the cotyledon epidermis and spontaneous cell death in the root meristem similar to what has been reported in another RBR-reduced line (Supplemental Fig. S1H, and Nowack et al., 2012; Borghi et al., 2010). These data support the previous observations that even slight modifications in its level influence RBR’s growth regulatory role. We tested the growth of seedlings of the above transgenic Arabidopsis lines having altered RBR levels and responses together with the WT control at a continuous temperature of 22°C, and after transferring them to 28°C 4DAG at 22°C (Fig. 2A). The root length of the seedlings was measured daily during the period of three more days (Fig. 2B). Root length of the three lines was comparable at 22°C. In comparison to the WT control roots, the *rbr1-2* line developed slightly shorter roots at 28°C, while strong repression was observed in the ectopic RBR-GFP line (Fig. 2A-B). To investigate why these transgenic RBR lines produced shorter roots than the WT control when grown at a higher temperature, we studied root meristems (RMs) after staining them with propidium iodide and analyzed them under confocal microscopy (Fig. 2C). The meristem size was calculated by counting the cortex cells from the quiescent centre (QC) to the transition zone, where they doubled in size (Fig. 2D). Three days after transferring the seedlings to 28°C (7 days in total after germination) the number of cortex cells dramatically increased in the root meristem of all three lines but at different levels (Fig. 2D). In the *rbr1-2* line, the number of cortex cells was found to be higher than in the WT at both 22°C and 28°C. In contrast, a much smaller number of cortex cells were counted in the ectopic RBR-GFP expressing roots at both 22°C and 28°C compared to the control WT. However, warm temperatures were also able to increase the number of cortex cells in this line, indicating that the cell proliferation inhibitory function of RBR may be partially suppressed by 28°C (Fig. 2D). We also determined the length of cortex cells in these genotypes at both temperatures. The cortex cells started to elongate in the RBR-GFP line much closer to the quiescent centre cells than in the WT, while the reduced RBR level in the *rbr1-2* mutant line had the opposite effect by delaying cell elongation in the root tip (Fig. 2E). We have observed a similar trend when growing these seedlings at a continuous temperature of 22°C and 28°C for either 7 or 14 days after germination (Supplemental Fig. S2). In all three lines, the root length increased over time, with the degree of increase dependent on both the temperature and the RBR level (Supplemental Fig. S2B,D). Accordingly, the warmer temperature likely activated cell proliferation in the RM of all investigated lines. However, the RBR level was inversely correlated with the number of cells in the root meristem at both temperatures. The number of cortex cells increased the most at 28°C in the *rbr1-2* mutant roots, while the increase was the lowest in the RBR-GFP roots two weeks after germination (Supplemental Fig. S2C, E). To further support that warm temperatures activate cell proliferation in the RM, we carried out a DNA replication-based assay. The seedlings were incubated with the thymidine analogue 5’-ethynyl-2’-deoxyuridine (5µM EdU) for 30 minutes to label the cells that are in the S-phase of the cell cycle. The roots of the three lines were labeled with EdU at 8, 24, or 48 hours after being transferred from 22°C to 28°C or remaining at 22°C for the same period. Interestingly, most of the S-phase cells were detected 8 hours after the seedlings were transferred to 28°C, indicating that entry into the cell cycle was quickly activated by the warm temperature in all three lines (Fig. 2F). However, as we have seen earlier in the case of the size of the root meristems, the EdU signal was negatively correlated with RBR levels, being strongest in the *rbr1-2* mutant and lowest in the RBR-GFP expressing line (Fig. 2G).

**Figure 2.**
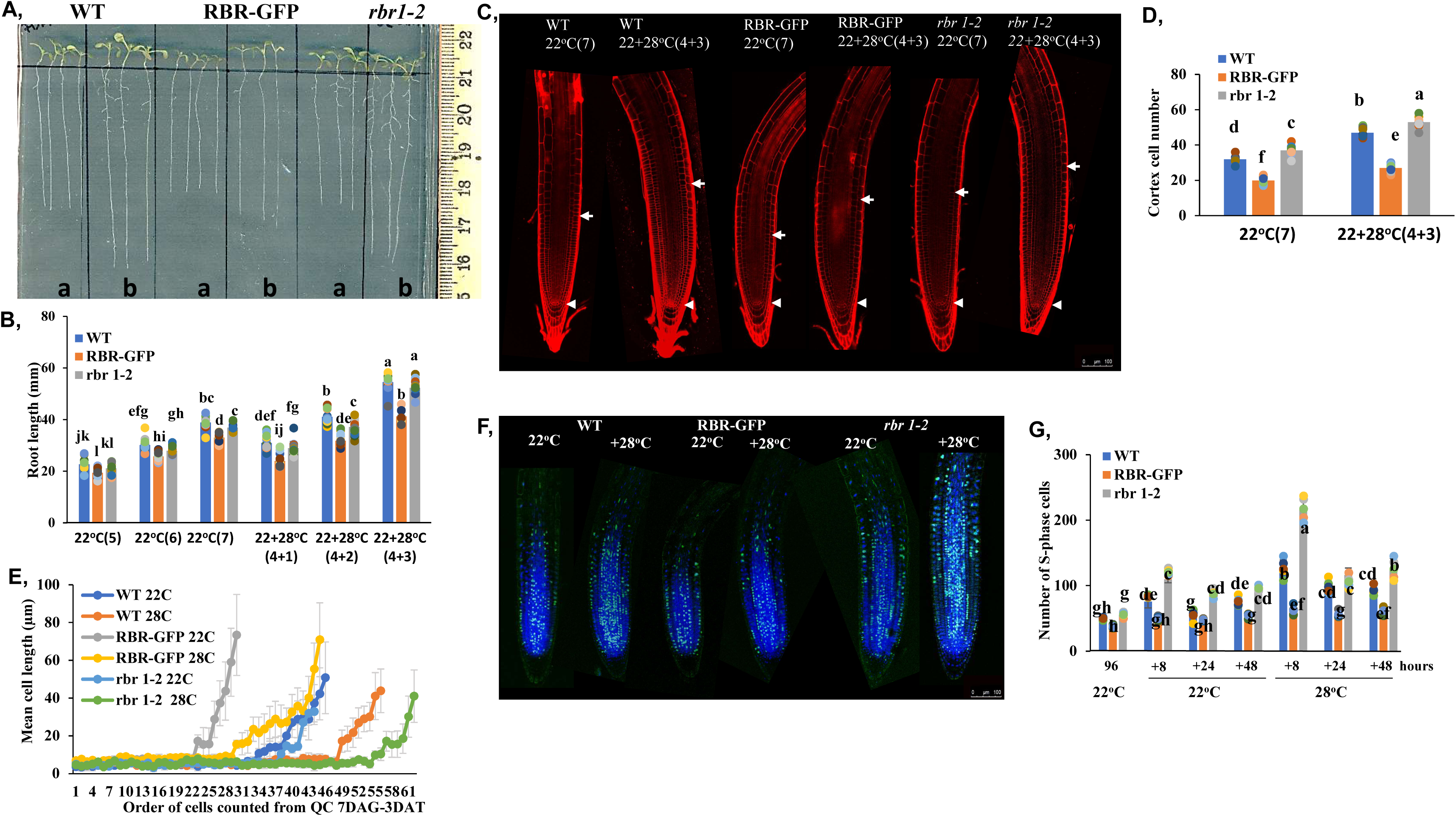
Ectopic RBR inhibits the warm temperature-induced root growth acceleration by repressing cell proliferation. Wilde type (WT), ectopic RBR-GFP expressing, and *rbr1-2* mutant seedlings were grown continuously at 22°C (control) or transferred from 22°C to 28°C (22+28°C) for the number of days indicated in the parentheses. **A**) Images of seedlings from the investigated lines grown on vertical plates at either 22°C(7) **(a)** or 22°C+28°C(4+3) **(b)**. The scale bar is shown on the right side. **B**) Comparison of root length increase between the 5th and 7th days in WT, RBR-GFP and *rbr1-2* seedlings grown continuously at 22°C or transferred on the fourth day to 28°C. n = 15. Different letters mean statistically significant differences (p < 0.05) based on one-way ANOVA analyses with Tukey’s HSD post-hoc test. **C**) Representative images of root tips of WT, RBR-GFP and *rbr1-2* seedlings grown at 22°C(7) for 22+28°C(4+3). The roots were stained with propidium iodide, and imaged under a confocal laser scanning microscope. Arrowheads indicate quiescent centers; arrows show the boundary of the root meristem in the cortex cell file. **D**) Cortex cell number in the root meristem of seven-day-old WT, RBR-GFP and *rbr1-2* seedlings grown at 22°C(7) or 22+28°C(4+3) (n=8 roots). Different letters mean statistically significant differences (p < 0.05) based on one-way ANOVA analysis with Tukey’s HSD post-hoc test. **E**) Spatial distribution of cell length across root meristems in WT, RBR-GFP and *rbr1-2* seedlings grown either at 22°C(7) or 22+28°C(4+3). Mean cell length was calculated at each position along the cortical cell file, which is defined by counts of cortical cells from the quiescent centre. Values are averages from the data analysed in eight roots from different plants (±SD). **F**) EdU (light blue-green) and Hoechst (blue) co-staining of nuclei in root tips of WT, RBR-GFP and *rbr1-2* lines grown at 22°C for four days and transferred to 28°C (+28°C) or left at 22°C for additional eight hours before a 30 min. incubation with 5 μM EDU labelling cells passing the S-phase of the cell cycle. **G)** The number of EdU-positive root cells was counted at the indicated time points from eight roots for each genotype at each time point. Data are average +/- standard deviation (n=3 biological replicates, N=8 samples in each). Different letters mean statistically significant differences (p < 0.05) based on one-way ANOVA analysis with Tukey’s HSD post-hoc test.

We also analyzed the shoot apex of young seedlings using scanning electron microscopy. As we have seen earlier in the case of WT seedlings, the leaf primordia of both RBR-GFP and *rbr1-2* lines were more developed just two days after being transferred from 22°C to 28°C (Supplemental Fig. S3A). However, we observed smaller and less developed leaf primordia in the shoot apex of RBR-GFP seedlings compared to the *rbr1-2* mutant and WT seedlings at both temperatures (Supplemental Fig. S3A). The average cell size in the 3^rd^ leaf primordium epidermis was comparable among these lines at both temperatures, indicating that increasing the RBR level decreases proliferation activity (Supplemental Fig. S3B). In response to the temperature transfer, the first leaves of all three lines developed faster and grew larger (Supplemental Fig. S3C,D and Fig. 3A). However, the *rbr1-2* mutant leaves reached a similar size as the wild-type ones, while the ectopic RBR-GFP leaves remained significantly smaller. By measuring the size of palisade cells, we could also estimate their numbers in these leaves (Fig. 3B,C,D). Warm temperatures increased both cell size and cell numbers in all three cases. However, while an elevated level of RBR had a negative effect on cell number, it had a positive effect on cell size. Conversely, the reduced level of RBR in the *rbr1-2* mutant leaf had a positive effect on cell number but negatively influenced cell size (Fig. 3C-D). Altogether, these data indicate that warm temperatures activate cell proliferation in the shoot apex and in young leaves similarly to the root tip and RBR is likely involved in this regulation. To further support this hypothesis, we analysed the expression of the cell cycle regulatory and E2F-RBR target genes, *ORC2* and *CDKB1;1* (Őszi et al., 2020; Gombos et al., 2023), in RBR-GFP and *rbr1-2* seedlings, as well as in the WT control soon after transferring them from 22°C to 28°C (Fig. 3E-F). Both the S-phase (*ORC2*) and the G2-M phase specific (*CDKB1;1*) regulatory genes were up-regulated in all three lines eight hours after the transfer. Afterwards, they both remained expressed at a higher level than at 22°C. The ectopic and reduced RBR-expressing lines showed opposite effects on the expression levels of these cell cycle genes increasing and decreasing them, respectively, in relation to the expression in the WT (Fig. 3E-F).

**Figure 3.**
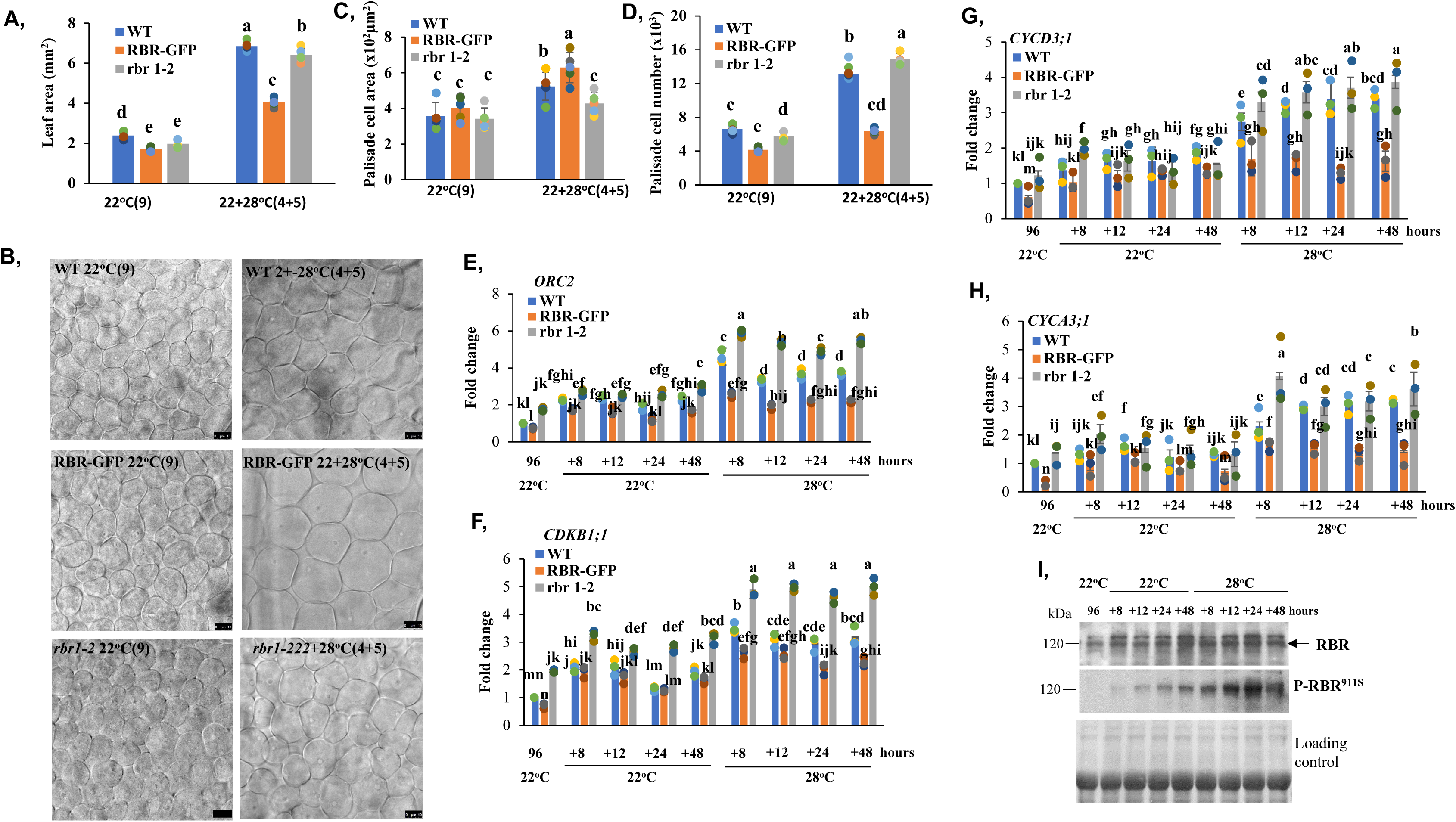
Warm temperatures suppress the cell proliferation inhibitor function of RBR through activating G1 cyclins and RBR-phosphorylation. Wilde type (WT), ectopic RBR-GFP expressing, and *rbr1-2* mutant seedlings were grown continuously at 22°C (control) or transferred from 22°C to 28°C (22+28°C) for the number of days indicated in the parentheses. **A-D**) Warm temperatures increase leaf size in an RBR-dependent manner. Size of the first leaf (A), images of palisade cells of the first leaf (**B**), the measured palisade cell area (n=5 leaf. ≥40 cells) (**C**), and the calculated cell number (leaf area/palisade cell area) (**D**). **E-F**) The S- and the G2-M-phase specific *ORC2* (**E**) and *CDKB1;1* (**F**) genes were induced by warm temperatures, depending on RBR. **G-H**) Transcript levels of the G1 cyclins, *CYCD3;1* and *CYCA3;1*, were elevated by warm temperatures, and are oppositely regulated in the RBR-GFP and *rbr1-2* lines. Expression of all the four genes (E-H) was monitored by qRT-PCR. Seedlings of WT, RBR-GFP and *rbr1-2* lines were grown for 96 hours at 22°C after germination and the samples were collected at the indicated time points at 22°C or 28°C, respectively. Values represent fold change normalized to the value of the relevant transcript of the seedlings at T0 (96 hours), which was set arbitrarily at 1. Data are means +/- sd. N=3 biological replicates. Different letters mean statistically significant differences (p < 0.05) based on one-way ANOVA analyses with Tukey’s HSD post-hoc test. **I**) Western blot shows RBR and phospho-RBR (P-RBR at the 911 serine site) levels in WT seedlings grown as indicated. Sampling was made as in E-H. The membrane was stained with comassie brilliant blue to indicate equal loading. Molecular weight marker (kDa) is shown at the left side. Arrow marks the position of the RBR protein.

RBR is known to be regulated by phosphorylation and we suggest that warm temperatures may repress RBR’s cell cycle inhibitory function by stimulating and increasing its phosphorylation. To investigate this hypothesis, we first studied the expression of two G1 *CYCLIN* genes: *CYCLIN D3;1* (*CYCD3;1*) and *CYCLIN A3;1* (*CYCA3;1*) (Fig. 3G,H). These genes are known to be the regulatory subunits of the plant RBR-kinase *CYCLIN-DEPENDENT KINASE A;1* (*CDKA;1 –* Van Leene et al., 2011; Takahashi et al., 2010). They both exhibited a similar expression pattern under our experimental conditions, similar to that of the cell cycle regulatory genes (Fig. 3E-H). They showed an early up-regulation at 8 hours of 28°C and afterwards maintained a generally and significantly higher expression level at 28°C compared to 22°C. Additionally, their expression levels were oppositely regulated by RBR; they were low in the ectopic RBR line, while high in the *rbr1-2* mutant indicating that they are E2F-RBR target genes (Bouyer et al., 2018; Őszi et al., 2020; Gombos et al., 2023; Fig. 3G-H). These data support the idea that RBR-kinase(s) could be activated by warm temperatures. To investigate this hypothesis, we monitored the phosphorylation of RBR through a western blot assay using a phosphosite-specific anti-RETINOBLASTOMA protein antibody. This antibody has been shown to label the Arabidopsis RBR protein in a serine 911 (911S) phosphorylation-dependent manner (Magyar et al., 2012; Wang et al., 2014). Previously, it was demonstrated that the RBR protein, phosphorylated on 911S, could not form a complex with E2Fs (Magyar et al., 2012). As shown in Fig. 3I, RBR became increasingly phosphorylated over time in WT seedlings at 22°C, indicating its developmental control. However, the level of the phosphorylated RBR form was much higher at 28°C, already 8 hours after the transfer, and continued to elevate afterwards.

All of this data supports the view that warm temperatures activate cell proliferation in young seedlings by suppressing the cell cycle inhibitor RBR through stimulating its phosphorylation.

### RBR positively regulates hypocotyl elongation induced by warm temperatures

Contrary to the apical meristem, the hypocotyl consists of mostly non-dividing post-mitotic cells. In the hypocotyl epidermis, there are only a few dividing cells, and they are all concentrated in the non-protruding longitudinal cell files belonging to the stomatal lineage in the upper half of the hypocotyl (Kono et al., 2007). Interestingly, the number of cells was previously found to be slightly elevated in the hypocotyl epidermis when the temperature was increased to 28°C, but cell elongation was considered to be the dominant process driving hypocotyl elongation at the higher temperature (Bellstaedt et al., 2019). Increasing the amount of RBR slowed down the acceleration of growth in both the shoot and root meristems under warm temperature (Figs 1 and 2). However, surprisingly, the hypocotyl of RBR-GFP expressing seedlings grew significantly longer after transfer from 22°C to 28°C compared to that of the WT (Fig. 4A,B). Contrary to the RBR-GFP expressing seedlings, the hypocotyl length of the *rbr1-2* mutant seedlings was shorter compared to the WT at 28°C (Fig. 4A-C). To understand the reasons for these phenotypic differences, we first measured the length of epidermal cells in the hypocotyl epidermis of WT, RBR-GFP and *rbr1-2* seedlings grown at either 22°C or after being transferred from 22°C to 28°C at 4 DAG. The length of the epidermal cells was found to be comparable in all of these seedlings grown at 22°C, but it started to differ soon after the seedlings were transferred to 28°C (Fig. 4C,D). Accordingly, epidermal cells elongated the most in the hypocotyl of the RBR-GFP seedlings, while they were less elongated in the *rbr1-2* mutant than in the WT (Fig. 4C,D and Supplemental Fig. S4A-C). The difference in cell elongation might be explained by differential ploidy increase in the hypocotyl cells in response to the elevated temperature. Although the ploidy level was indeed increased in the hypocotyl of all the three lines at warm temperatures as more 16C nuclei were detected at 28°C than at 22°C, the increase was at a comparable level in all the three lines (Supplemental Fig. S4D). In the WT, the number of cells in the hypocotyl also increased in response to the elevated temperature, confirming previous observations (Bellstaedt et al., 2019 and Supplemental Fig. S5). The *rbr1-2* mutant had more cells but showed a similar temperature response than the WT (Supplemental Fig. S5). The number of hypocotyl cells also increased in the RBR-GFP line, but the cell number and the increase were lower in comparison to the other two lines (Supplemental Fig. S5). Interestingly, small meristemoid-like cells belonging to the stomatal lineage were found to be the most proliferative in the *rbr1-2* mutant, and warm temperatures further increased their number (Supplemental Fig. S6). In contrast, ectopic RBR reduced the numbers of these cells at both temperatures compared to the WT (Supplemental Fig. S6). These data indicate that the amount of RBR positively correlates with cell length but inversely with cell number in the hypocotyl. Previously, we have shown that canonical E2Fs are the primary effectors of RBR in Arabidopsis and the three E2Fs redundantly bind to a similar set of genes involved in the control of cell proliferation, together with RBR (Gombos et al., 2023). To test whether RBR controls hypocotyl elongation through E2Fs, we introduced RBR-GFP into an E2FA mutant background by crossing the RBR-GFP line with a T-DNA insertion E2FA mutant line (*e2fa-1* – Leviczky et al., 2019). The mutation of E2FA did not change hypocotyl length in the presence of ectopic RBR-GFP, suggesting that RBR might regulate hypocotyl elongation independently of E2FA (Supplemental Fig. S7A-B).

**Figure 4.**
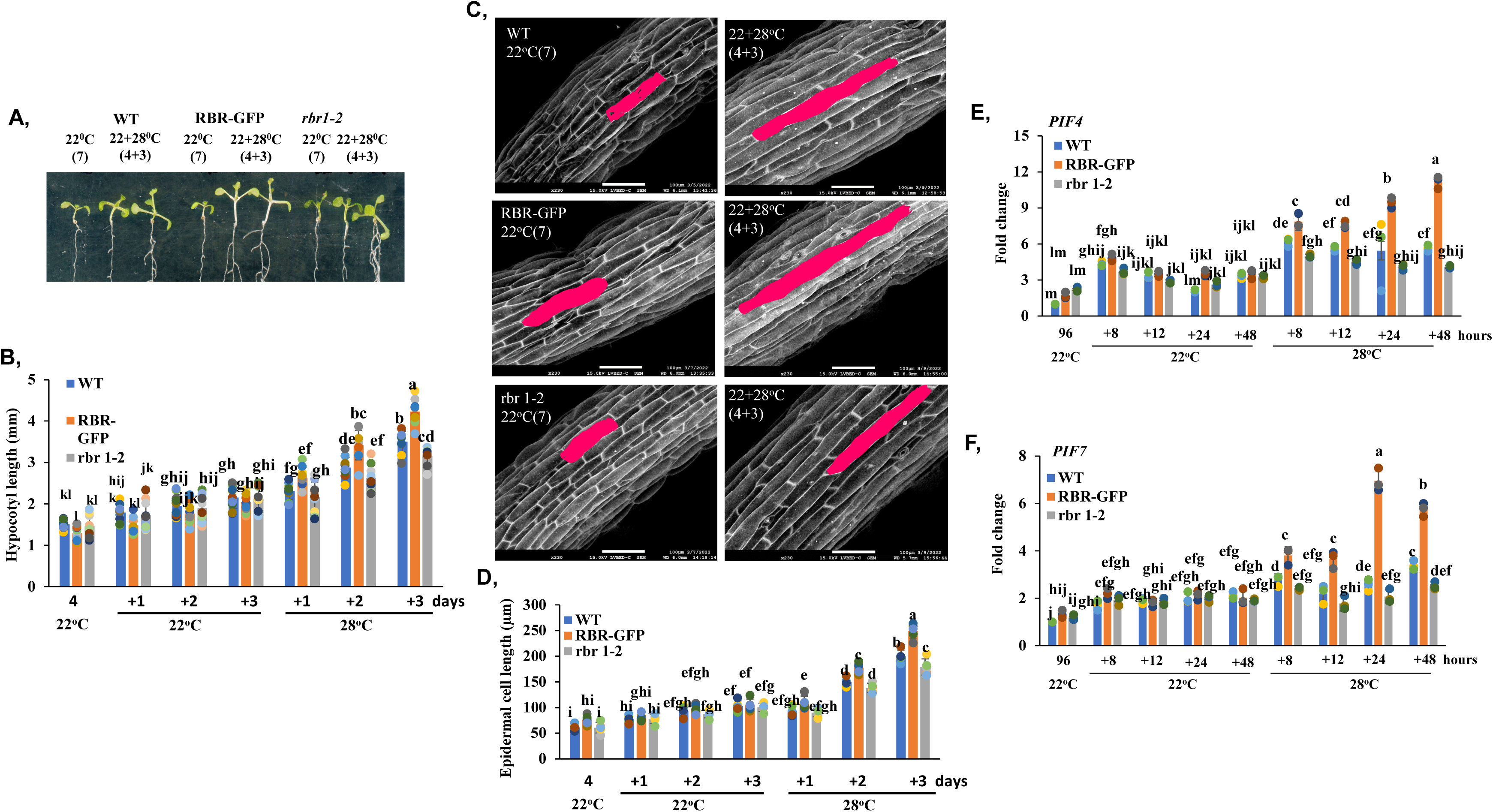
At warm temperatures, RBR stimulates the elongation growth of the hypocotyl, inducing the expression of PIF4 and PIF7 transcription factors. Wild type (WT), ectopic RBR-GFP expressing, and *rbr1-2* mutant seedlings were grown continuously at 22°C (control) or transferred from 22°C to 28°C (22+28°C) for the number of days indicated in the parentheses. **A**) Ectopic RBR increases hypocotyl length following the transfer of seedlings to 28°C. **B**) Hypocotyl length of the three genotypes with different RBR levels was measured both at continuous 22°C or after transfer to 28°C at the indicated time points. N=10; Different letters mean statistically significant differences (p < 0.05) based on one-way ANOVA analyses with Tukey’s HSD post-hoc test. **C**) Hypocotyl epidermis was analysed under a scanning electron microscope at both temperature regimes at the indicated time points. Scale bar is 100 µm. **D**) Cell length of hypocotyl epidermis in the three lines was measurred by using the ImageJ softwere. N=5. Different letters mean statistically significant differences (p < 0.05) based on one-way ANOVA analyses with Tukey’s HSD post-hoc test. **E-F**) Expression of *PIF4* (**E**) and *PIF7* (**F**) in WT, RBR-GFP and *rbr1-2* seedlings at the indicated temperatures and times. T0 represents 96-hour-old seedlings grown at 22°C. These were either kept growing at 22°C or after transfered to 28°C, and samples were taken at 8, 12, 24 and 48 hours afterwards. Values represent fold change normalized to the value of the relevant transcript of the seedlings at T0 (96 hours), which was set arbitrarily at 1. Data are means +/- sd. N=3 biological replicates. Different letters mean statistically significant differences (p < 0.05) based on one-way ANOVA analyses with Tukey’s HSD post-hoc test.

Cell elongation is stimulated in the hypocotyl by warmth through the PIF4/PIF7-YUCCA auxin regulatory module (Casal & Balasubramanian, 2019). We were curious about whether RBR influences the expression of the main regulators of thermomorphogenesis. For this purpose, we grew seedlings for 4 days at 22°C. Afterwards, we either continued their culture at 22°C or transferred them to 28°C. We conducted a q-RT-PCR experiment to determine the expression of *PIF4* and *PIF7*, at 8, 24, and 48 hours after transfer. As shown in Figure 4 E, F, both genes were expressed at comparable levels in all three lines at 22°C. However, this picture dramatically changed when transferring the seedlings to 28°C. Both *PIF* genes were already upregulated in each line just 8 hours after warm treatment. The strongest activation was found in the ectopic RBR expressing line, while *rbr1-2* was less effective in the upregulation (Fig. 4E,F). In addition, *PIF4/7* expression continuously increased in the RBR-GFP line at 28°C, much more than their expressions change in the WT and in the *rbr1-2* mutant line at 28°C during the investigated period (Fig. 4E,F). Emerging data support the view that PIFs activate the expression of auxin biosynthetic *YUCCA* genes under shade and warm temperatures, resulting in auxin-mediated elongation growth (Casal & Balasubramanian, 2019). To test whether ectopic RBR could also stimulate the expression of *YUCCA* genes, we analyzed the expression of four *YUCCA* genes (*YUC1,2,8*, and *9*) including *YUC8* a rate-limiting auxin biosynthesis enzyme in temperature-dependent hypocotyl elongation in the same experiments described above (Fig. 5A-D). Interestingly, all these *YUCCA* genes showed similar expression patterns to the *PIF4* and *7* genes, as they were all up-regulated by warm temperatures in all three lines 8 hours after the seedlings were transferred to 28°C. Additionally, their expression levels correlated well with RBR levels, with the highest expression values in the ectopic RBR line and the lowest ones in the *rbr1-2* mutant line (Fig. 5). In addition, *TRANSPORT INHIBITOR RESPONSE 1* (*TIR1*), an auxin receptor F-box protein expressed similarly to *PIF4-7* in these lines at warm temperatures (Supplemental Fig. S8A). These data further support that RBR could accelerate hypocotyl elongation at 28°C by positively regulating the PIF-YUCCA regulatory module and auxin signaling.

**Figure 5.**
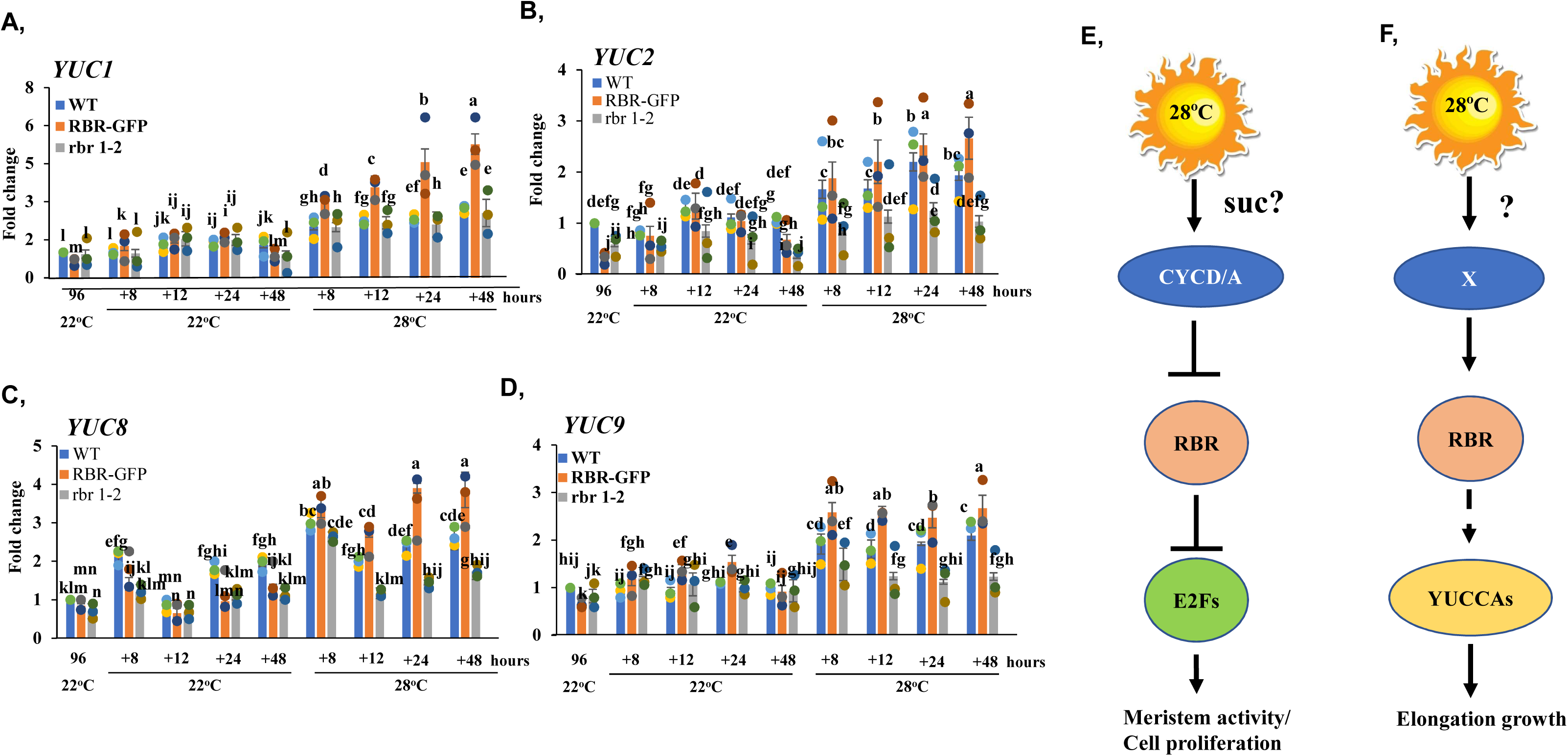
RBR positively regulates the auxin synthesis regulatory *YUCCA* genes. **A-D**) Expression of four *YUCCA* genes were followed by qRT-PCR as indicated. T0 represents 96-hour-old seedlings, and they were either kept growing at 22°C or transferred to 28 °C, and the samples were taken at 8, 12, 24 and 48 hours afterwards. Values represent fold change normalized to the value of the relevant transcript of the seedlings at T0 (96 hours), which was set arbitrarily at 1. Data are means +/- sd. N=3 biological replicates. Different letters mean statistically significant differences (p < 0.05) based on one-way ANOVA analyses with Tukey’s HSD post-hoc test. **E-F**) Schematic model how temperature regulates cell proliferation and elongation growth through modifying RBR activity. **E**) Warm temperatures activate G1 cyclins like CYCD3;1 and CYCA3;1. This might be mediated by the increased sucrose level due to the temperature-enhanced metabolic activity. The cell cycle inhibitor RBR is phosphorylated and repressed by an RBR-kinase complex, such as CDKA;1-CYCD/A. This results in RBR-free E2Fs, which drive cell cycle entry and accelerate meristematic function in both the root and shoot apices. **F)** Warm temperatures might cause post-translational modifications on RBR, such as phosphorylation at unknown site(s) in post-mitotic cells. This could result in the stimulation, either directly or indirectly, of the expression of non-canonical targets, like YUCCAs.

In contrast to *PIF4-7*, the expression of *ELONGATED HYPOCOTYL 5* (*HY5)*, a key regulatory gene in photomorphogenesis, was similar to that of cell cycle genes under warm temperatures (Supplemental Fig. S8B and Fig. 3). Its level increased in the *rbr1-2* mutant, while it decreased in the RBR-GFP line compared to the WT. Accordingly, RBR exerts opposite control over the expression of *HY5* and *PIF4-7*, the regulators that play antagonistic roles in both light and temperature-induced developmental processes (Wang and Zhu, 2022).

## Discussion

Plants exposed to non-stressful warm temperatures change their growth rate and accelerate their development. It is suggested that plant thermo-responses could either be passive or active and growth acceleration was speculated to be a general response caused by thermodynamic effects on metabolic rates and enzyme activities (Casal and Balasubramanian, 2019). In favour of this theory, recently ten Arabidopsis accessions were analysed at different temperatures and all showed similar robust growth accelerations throughout the vegetative phase under warm temperatures indicating that genotype has just limited effect on this temperature induced growth-related phenotype (Ibanez et al., 2017). All of these accessions, however, might have intact and conserved signalling elements responsible for robust temperature-induced growth responses.

Here we show that warm temperature activates cell proliferation in the meristems both in the shoot and in the root apices where cell division in plants is concentrated. We demonstrate that cell proliferation in Arabidopsis seedlings is rapidly activated soon after temperature was raised. A recent study has already pointed to cell proliferation as a major process to accelerate root growth (Ai et al., 2023). Here, we not only confirm their findings, but also show that leaf primordia in the shoot apex and the developing young leaves grow faster under warm temperatures. This is because they produce more cells than at normal growth temperatures. We suggest that warm temperatures accelerate meristem functions through the conserved CYCD-RBR-E2F molecular pathway. This is supported by the following evidence: *(i)* the expression of G1 cyclins such as *CYCD3;1* and *CYCA3;1* rapidly increased at elevated temperatures (Fig. 3G,H), *(ii)* at the same time, the phosphorylation of the cell cycle inhibitor RBR at a conserved CDK site was augmented (Fig. 3I), and *(iii)* the expressions of E2F-target cell cycle regulatory genes *ORC2* and *CDKB1;1* were also up-regulated with warm temperatures (Fig. 1M,N; Fig. 3E,F).

How warm temperatures could activate the canonical cell cycle regulatory CYCD-RBR-E2F pathway? Plant cyclin Ds have already been implicated to mediate both internal and environmental signals toward the cell cycle control system (DeVeylder, 2019; Dewitte et al., 2007). Interestingly, warm temperatures induce a similar growth acceleration phenotype in transgenic plants overexpressing CYCD2;1 without affecting the final organ size and plant stature (Cockroft et al., 2000). In the case of ectopic CYCD2;1, it was shown that cells enter the cell cycle faster by shortening the G1 phase (Cockcroft et al., 2000). It is also very possible that warm temperatures similarly affect the length of the cell cycle. The increased number of S-phase cells in the root meristem, just eight hours after the temperature was shifted, supports the idea that cells are entering the cell cycle at a faster rate.

Expression of Arabidopsis *CYCD2;1* and *CYCD3;1* both quickly respond to the sucrose level (within 30 minutes and 4 hours, respectively) in the nutrient medium (Riou-Khamlichi et al., 2000). Overexpression of CYCD3;1 activates mitosis and stimulates the phosphorylation of RBR even in nutrient limited condition (Magyar et al., 2012). We show here that warm temperature also rapidly activates the expression of *CYCD3;1* indicating that CYCDs might be targeted soon after temperature was sensed. It is tempting to speculate that warm temperatures activate *CYCD3;1* (and/or *CYCD2*) by changing nutrient and sucrose levels, due to increased metabolic rates in the plants. Recently it was noted that the expression of genes involved in sucrose transport and response in the root, and shoot, respectively, was quickly upregulated (within 4 hours) by warm temperature (Gaillochet et al., 2020).

According to the classical plant cell cycle model, CYCDs activate the RBR-kinase CDKA;1, and hyperphosphorylation of RBR represses its cell cycle inhibitory function. RBR has numerous conserved CDK sites but only few has been experimentally verified in plants (Desvoyes and Gutierrez, 2020). Previously, it has been shown that phosphorylation of RBR at position 911, on the serine residue of a conserved CDK site, represses its ability to bind to E2Fs (Magyar et al., 2012). In addition, phosphorylation of this site was observed in dividing cells and tissues, and its level increases when cells enter the cell cycle (Ábrahám et al., 2011; Magyar et al., 2012). Therefore, the phosphorylation of RBR at this site indicates a higher proliferation activity in seedlings growing at warmer temperatures (Fig. 3I). Our data support that warm temperatures act on meristems as a mitogenic factor, and its signal is mediated through the canonical CYCD-RBR-E2F pathway, resulting in faster growth and accelerated development (Fig. 5E).

Unexpectedly, RBR not only functions as a cell cycle inhibitor, but also exerts a positive influence on the elongation growth of post-mitotic, quiescent cells under warm conditions. It is quite remarkable that the hypocotyl of the ectopic RBR-GFP expressing line grows significantly longer than that of the WT at warm temperatures (Fig. 4.A-E). The RBR protein could be detected in epidermal cells of the hypocotyl at both temperatures, regardless of whether they were proliferating or quiescent, supporting its regulatory role in both cell types (Suplemental Fig. S1D). In the *rbr1-2* mutant, proliferation of meristemoid cells of the stomatal lineage were activated supporting the canonical cell cycle regulatory role of RBR in these cells, the division of which was further stimulated by warm temperatures indicating that elevated temperature has a positive effect on cell proliferation even in the hypocotyl confirming previous findings (Suplemental Fig. S1E and S6B, Bellstaedt et al., 2019).

The hypocotyl of the RBR-GFP expressing line grows longer at warm temperatures because cell elongation is stimulated by the ectopic RBR. This effect appears to be specific, as epidermal cells in the hypocotyl of the *rbr1-2* mutant line were less elongated compared to the WT control. This suggests that RBR regulates the cell size of post-mitotic hypocotyl cells under warm temperatures. The animal Rb protein and its functional budding yeast analogue, Whi5, function as cell size controllers during cell proliferation (Zatulovskiy et al., 2020; Schmoller et al., 2015). However, different cell cycle inhibitor molecules have been proposed to operate in dividing plant cells (D’Ario et al., 2021). How can the cell cycle inhibitor RBR stimulate elongation growth in post-mitotic hypocotyl cells?

Surprisingly, thermo-morphogenic regulatory genes including the central *PIF4* and *PIF7* together with their target auxin biosynthetic *YUCCA* genes (*YUC1-2* and *YUC8-9*) were all found to be positively regulated by RBR at the seedling level (Fig. 4E,F; Fig. 5A-D). This suggests that RBR mediates the effect of warm temperatures on hypocotyl elongation as a direct or indirect activator of gene expression. In dividing cells, elevated expression of CYCDs drives the RBR kinase, resulting in an activated cell cycle. However, we do not yet know what signal is present in elongated post-mitotic hypocotyl cells that leads to RBR activation. Interestingly, CYCDs such as CYCD3;1 are expressed at the highest levels in the hypocotyl, suggesting that CYCD3;1 may also have a function in non-dividing cells similar to that of RBR. In animals, it has been shown that monophosphorylated forms of Rb are present in various protein complexes and they function beyond the regulation of the cell cycle (Sanidas et al., 2019).

RBR may control the expression of thermo-regulatory genes, either directly or indirectly. RBR itself cannot bind to target gene sequences, but it regulates cell cycle control genes in complex with its primary effector, E2F transcription factors (Gombos et al., 2023). Neither PIF4/7 nor YUCCA genes, however, contain conserved E2F-binding sequences in their regulatory promoter regions. In addition, RBR and E2Fs were not enriched in the promoter regions of these genes at normal growth temperature further supporting that they are regulated by RBR independently of E2Fs (Bouyer et al., 2018; Gombos et al., 2023). Additionally, hypocotyl elongation stimulated by ectopic RBR at warm temperature was not affected by mutating E2FA, one of the three canonical E2Fs (Supplemental Fig. S7), however we cannot exclude that RBR might work together with any of the other two E2Fs under warm temperatures.

Altogether, our data indicated that warm temperatures have a dual effect on RBR activity. It suppresses its canonical cell cycle inhibitor function in both shoot and root meristems, driving faster growth and development. Additionally, it activates its non-canonical cell elongation regulatory function in the hypocotyl by increasing, among others, the expression of auxin biosynthetic YUCCA genes. In both pathways, CyclinDs might mediate the temperature control on RBR function, however, this hypothesis needs further experimental verification.

## Materials and Methods

### Plant material and growth conditions

Arabidopsis thaliana Col-0 ecotype was used as WT control, and was the background of the other transgenic lines utilized in this study. Most of the transgenic lines have been previously published: RBR-GFP (Magyar et al., 2012), CYCB1;2-YFP (Iwata et al., 2011), *rbr1-2* mutant (Nowack et al., 2006), *e2fa-1* mutant (Leviczky et al., 2019). We generated an *e2fa-1*/RBR-GFP double transgenic line by crossing *e2fa-1* mutant line with the RBR-GFP expressing line.

Seeds were surface sterilized, rinsed with sterile water and subsequently stratified for 72 hours in the dark (4°C). After sowing, seeds were germinated and cultivated on half-strength germination medium supplemented with 1% (w/v) sucrose either on vertically or horizontally oriented plates in climate controlled growth cabinets (MLR-350, Sanyo, Gallenkamp, UK) at constant temperatures of 22°C or 28°C under long day conditions (16 h light/8 h dark) and with 100-120 µmol m^-2^ s^-1^ photosynthetically active radiation (PAR) from white fluorescent lamps. For temperature shift experiments, plants were grown for 4 days after germination (DAG) at 22°C and then half of them were kept on the same temperature while the other half were transferred to 28°C and grown for few more days afterward.

### Microscopy

For analysing root under confocal laser microscopy (SP5, Leica) seedlings were grown on vertically oriented plates, and roots were stained with propidium iodide (PI – 20 µg/mL) and photographed afterwards. Cell length were measured by using Image J software.

#### Microscopic observation of the first leaf

Young seedlings were fixed in a 9:1 of ethanol and acetic acid solution and cleared with Hoyer’s solution (a mixture of 100 g chloral hydrate, 10 g glycerol, 15 g gum arabic, and 25 mL water), we performed microscopic observations using the 1st leaf. After whole leaf images were captured, palisade cells at positions one-fourth and three-fourth from the tip of each leaf were observed with differential interference contrast (DIC) microscope (Leica). The captured images were analyzed using ImageJ (ver.2.1.0; rsb.info.nih.gov/ij) and the average size of palisade cell, the number of cells in the uppermost layer of palisade tissue per leaf, were calculated according to methods described previously (Nomoto et al., 2022).

#### Scanning electron microscopy observation of leaf primordium and hypocotyls

Young seedlings were vacuum infiltrated and fixed with 100% methanol for 20 min, dehydrated in 100% ethanol for 30 min and then in fresh 100% ethanol overnight. Next day samples were critical point dried, mounted on SEM stubs and observed in a JEOL JSM-7100F/LV scanning electron microscope in low-vacuum mode. High contrast cell outlines in the uncoated plants were imaged according to Talbot and White (2013) by detecting backscattered electrons at 15 kV accelerating voltage and 40 Pa pressure in the specimen chamber.

#### EdU staining of root meristem

For EdU incorporation assay WT and transgenic seedlings were grown at 22°C for 4 DAG, and half of the seedlings were transferred to 28°C or kept on 22°C for 8, 24, 48 hours when they were placed into half strength liquid MS containing 5 µM EdU (Click-iT Alexa Fluor 647 Imaging Kit; Invitrogene) and incubated for 30 minutes at the same temperatures. EdU detection was carried out according to Kazda et al., 2016. The root samples were also stained with HCS NuclearMask Blue stain provided in the kit (Invitrogene), and the observations were carried out under the confocal laser microscope.

### RT-qPCR

RNA was extracted from young seedlings using a CTAB-LiCl method described as Jaakola et al., 2001. RNA samples were treated with DNase1 (ThermoScientific #EN0521) according to the manufacturer’s protocol. cDNA was synthesised using 1 μg of RNA using the ThermoScientific Reverse Transcription Kit (#K1691) with random hexamers based on the manufacturer’s prescription. Mock reaction without RevertAid enzyme was also prepared to ensure that there is no contaminating genomic DNA in the samples. RT-qPCR in the presence of SYBR Green (TaKaRa TB Green Primer Ex TaqII, #RR820Q) was carried out according to the manufacturer’s instructions in a BioRAD CFX 384 Thermal Cycler (BioRAD) with the following setup: 50 °C 2 min, 95 °C 10 min, 95 °C 15 s, 60 °C 1 min, 40 cycles followed by melting point analysis. Each reaction was carried out in tree technical replicates and reaction specificity was confirmed by the presence of a single peak in the melting curve. All the data were normalised to the average Ct value of two housekeeping genes (ACTIN and UBIQUITIN) unless otherwise mentioned and the calculated efficiency was added to the analysis. Amplification efficiencies were derived from the slope of amplification curves at the exponential phase. Primer sequences are summarised in Supplementary Table 1.

### Immunoblotting

For immunoblotting assay, we used whole seedlings, and proteins were extracted in extraction buffer (25mM Tris-HCl, pH 7.5, 75mM NaCl, 15 mM MgCl_2_, 15 mM EGTA, 15 mM p-nitrophenylphosphate, 60 mM β-glycerophosphate, 1mM dithiothreitol, 0.1% (v/v), IGEPAL CA-630, 0.5mM NaF, 1 mM phenylmethylsulfonyl fluoride, and protease inhibitor cocktail for plant tissue (P9599, Sigma). 40 µg extracted proteins were loaded on SDS-PAGE gel (8%), and after the gel electrophoresis proteins were transferred onto polyvinylidene difluoride membrane (PVDF, Milipore). Primary antibodies used in the immunoblotting experiments were chicken anti-RBR antibody (1:2000 dilution; Agrisera), and anti-phospho-specific Rb (Ser-807/811) rabbit polyclonal antibody (1:500 dilution; Cell Signalling Tech). Membrane was blocked in 5% (w/v) milk powder with 0.05% (v/v) Tween 20 in Tris-buffered saline (TBS; 25 mM tris-HCl, pH 8.0, and 150 mM NaCl; TBS plus Tween 20 TBST). Membrane was incubated with primary antibody for overnight on a shaker at 4°C, an after washing with TBST, next the membrane was incubated with the appropriate secondary antibody conjugated with horseradish peroxidase at room temperature. Chemiluminescence substrates were applied either purchased from Thermo Fisher Scientific (SuperSignal West Pico Plus) or from Milipore (Immobilon western horseradish peroxidase).

### Flow cytometry analysis of hypocotyls

For flow cytometry measurement, the hypocotyls were collected and chopped by razor blades in nuclei extraction buffer and stained with DAPI with the CyStain UV Precise Kit (Partec, Magyar et al., 2005). Nuclear DNA content was determined by using Partec PAS2 Particle Analysing system (Partec, Germany).

### Statistical assays

Quantitative data are presented as the mean ± SD and the statistical analysis between different groups was analysed by one-way analysis of variance (ANOVA) following Tukey’s HSD post hoc test to determine, which group are significantly different from each other; *p*-values of less than 0.05 were considered significant.

## Acknowledgements

We thank Gábor Steinbach (HUN-REN Biological Research Centre, Szeged, Hungary) for helping with microscopy. This work was supported by a grant from the Hungarian National Research Funding (NKFI-132486). Rasik Shiekh Bin Hamid was supported by the stipendium hungaricum fellowship.

## Supplementary

**Supplemental Figure S1.**
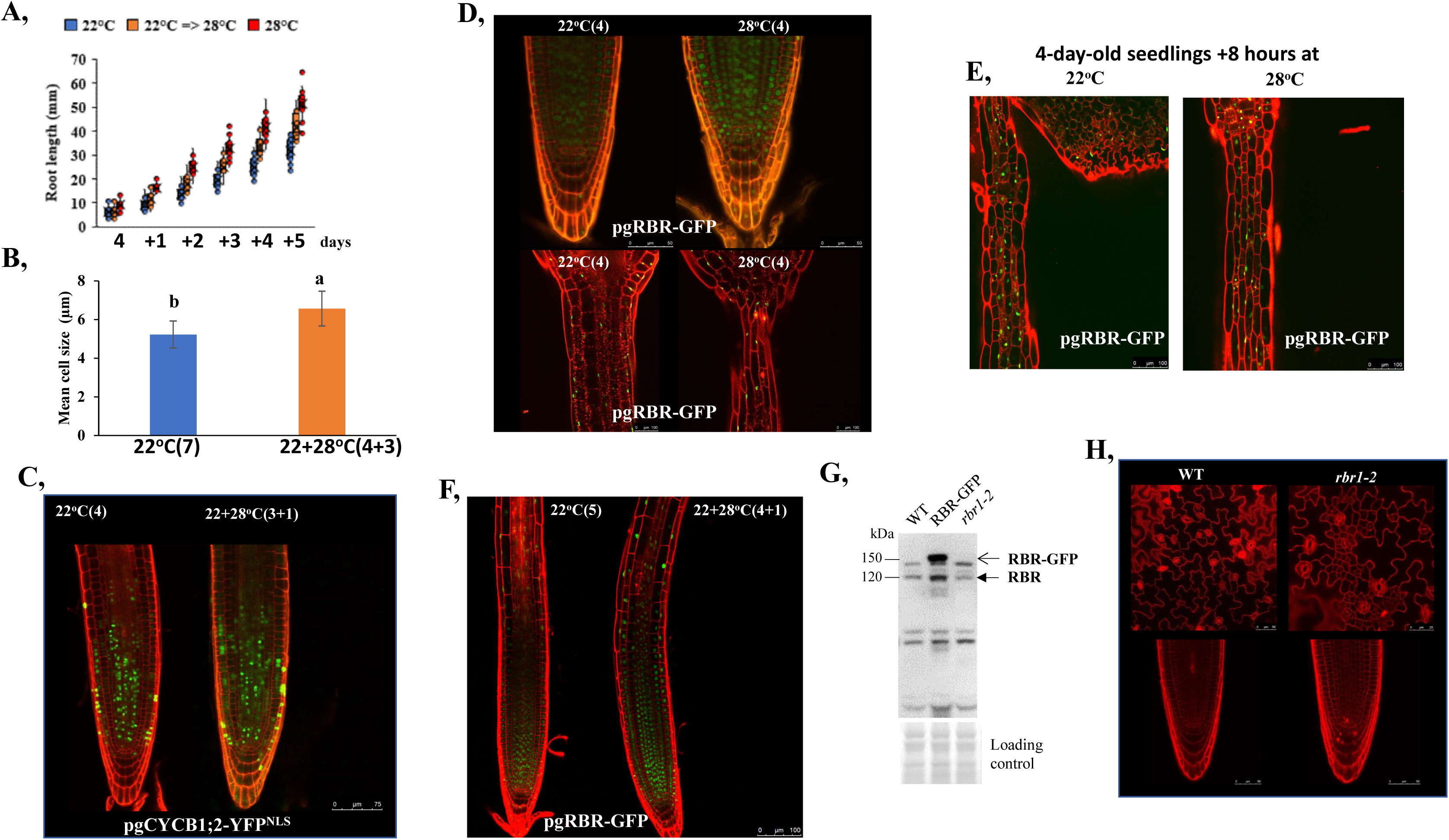
The RBR proteins accumulate in both mitotic and post-mitotic cells, and their amount is increased by warm temperatures. **S**eedlings were grown continuously at 22°C (control) or transferred from 22°C to 28°C (22+28°C) for the number of days indicated in the parentheses. **A**) Root length of wild-type (WT) seedlings at 22°C, 28°C, or after transfer from 22°C to 28°Cat the fourth day. The measuring started four days after germination. N=10. Three biological replicates. **B**) Mean cortex cell size in the root meristem of WT seedlings grown with or without transfering them from 22°C to 28°C. Values are averages from the data obtained by the analysis of eight roots from different plants (±SD). N=3 biological replicates. Different letters mean statistically significant differences (p < 0.05) based on one-way ANOVA analyses with Tukey’s HSD post-hoc test. **C**) Fluorescent images of propidium iodide-stained (red) roots expressing pgCYCB1;2-YFP^NLS^ (green). The roots are from seedling grown at 22°C for 4 days, or 3 days at 22°C plus one day at 28°C. Scale bar is 75 µm. **D**) Fluorescent images of propidium iodide-stained (red) root tips (upper row) and hypocotyls (bottom row) of seedlings expressing RBR-GFP (green) under the control the *RBR* gene promoter (pgRBR-GFP). The seedlings were grown at the indicated temperatures for four days after germination. **E,** The pgRBR-GFP signal is comparable in the hypocotyl epidermal cells of seedlings growing continuously at 22°C for four days plus eight hours, or transferred to 28°C for 8 hours after four days at 22°C. **F**) The GFP signal was increased at warm temperatures in the root tips of seedlings expressing pgRBR-GFP. **G**) Western blot analysis of RBR levels in WT, RBR-GFP and *rbr1-2* seedlings grown at long-day condition (16/8 hours light-dark cycle). The open arrow shows RBR-GFP, the black arrow marks the endogenous RBR protein. Ponceau-S-stained membrane was used as loading control (below). Molecular weight is marked at the left side. **H**) The cell proliferation and cell death rates are augmented in *rbr1-2* mutant cotyledon (upper pictures) and root tip (bottom images) in comparison to the similar-aged WT.

**Supplemental Figure S2.**
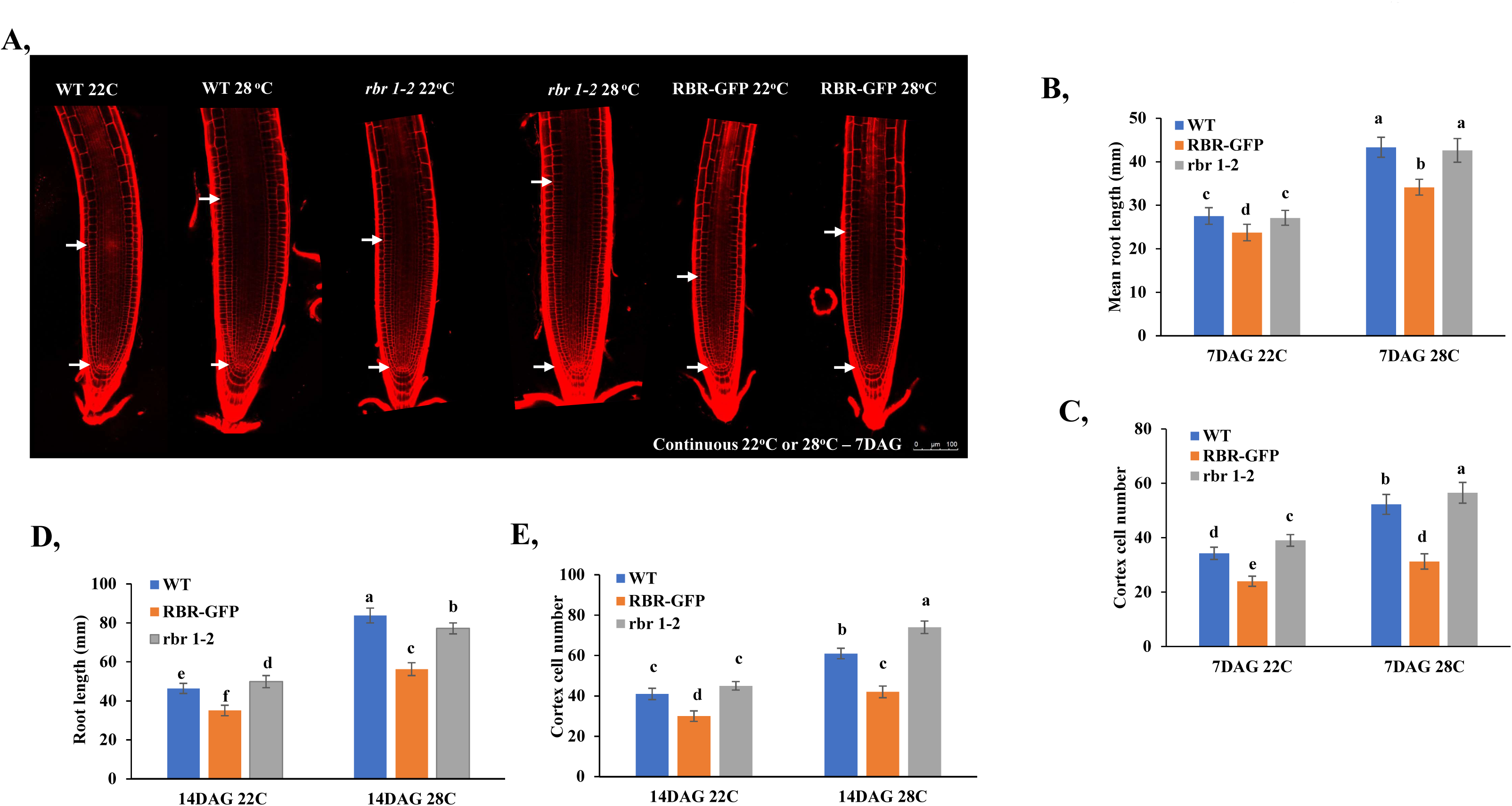
Elevated temperatures enlarge the size of the root meristem, a phenomenon that is modulated by varying levels of RBR. Wild type (WT), ectopic RBR-GFP expressing, and *rbr1-2* mutant seedlings were grown continuously at 22°C or 28°C, respectively, for the number of days after germination (DAG) as indicated. **A**) Representative images of root meristems of seven-day-old seedlings grown continuously at 22°C or 28°C. Arrows mark the quiescent centres and the last isodiametric small cortex cells, indicating the boundaries of the root meristem. **B**) Mean root length of the three genotypes at conditions as in A (n=15). **C**) Cortex cell numbers in the root meristems of the same lines at both temperatures (n=8). **D**) Mean root length was also determined in two-week-old seedlings of these genotypes at both temperatures (n=15). **E**) Cortex cell numbers in the same two-week-old seedlings at 22°C or 28°C (n=8) Different letters mean statistically significant differences (p < 0.05) based on one-way ANOVA analyses with Tukey’s HSD post-hoc test (B-E).

**Supplemental Figure S3.**
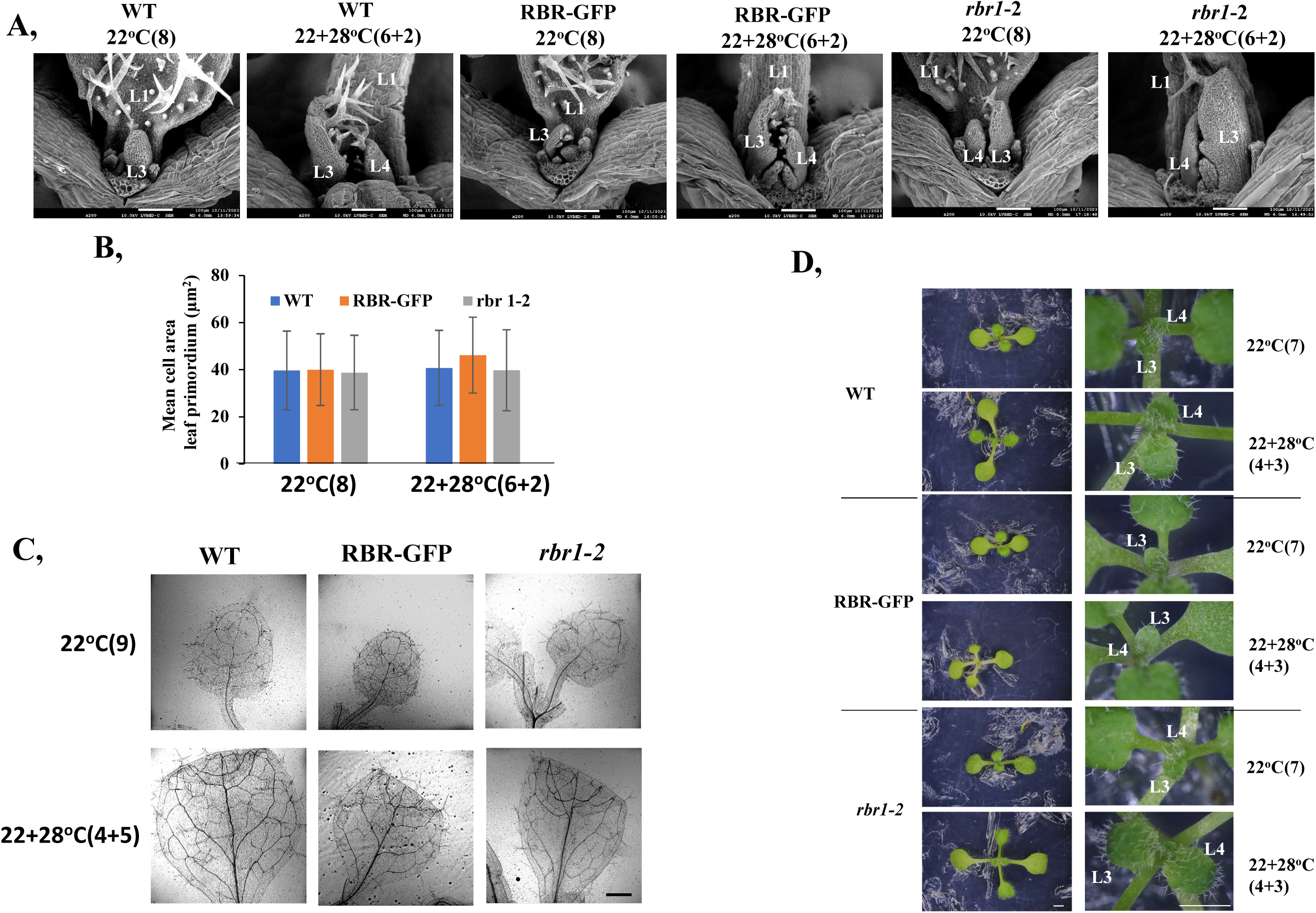
Warm temperatures accelerate shoot and leaf development and this effect is dependent on the RBR level. Wild type (WT), ectopic RBR-GFP expressing, and *rbr1-2* mutant seedlings were grown continuously at 22°C (control) or transferred from 22°C to 28°C (22+28°C) for the number of days indicated in the parentheses. **A**) Scanning electron microscopic images of shoot apices of eight-day-old wild type (WT), RBR-GFP and *rbr1-2* seedlings. Leaf primordium size increased in all the three genotypes following transfer from 22°C to 28°C at the sixth day. The order of leaf primordia (L) is marked by numbers. Scale bar is 100µm. **B**) Cell size of the third leaf primordium was measured in all the three lines at the indicated temperature regimes (n=8 with 45 or more cells were measured). **C**) The first leaf of the seedlings were imaged under a confocal laser scanning microscope at conditions as indicated. Scale bar is: 500 µm. **D**) Leaf primordium development was also investigated under a stereo microscope in all lines at the indicated conditions. Warm temperatures increased leaf primordium size in all the three lines, but the strongest effect was seen in the *rbr1-2* mutant, while ectopic RBR partially suppressed the temperature-induced growth acceleration. Scale bar is: 3 mm.

**Supplemental Figure S4.**
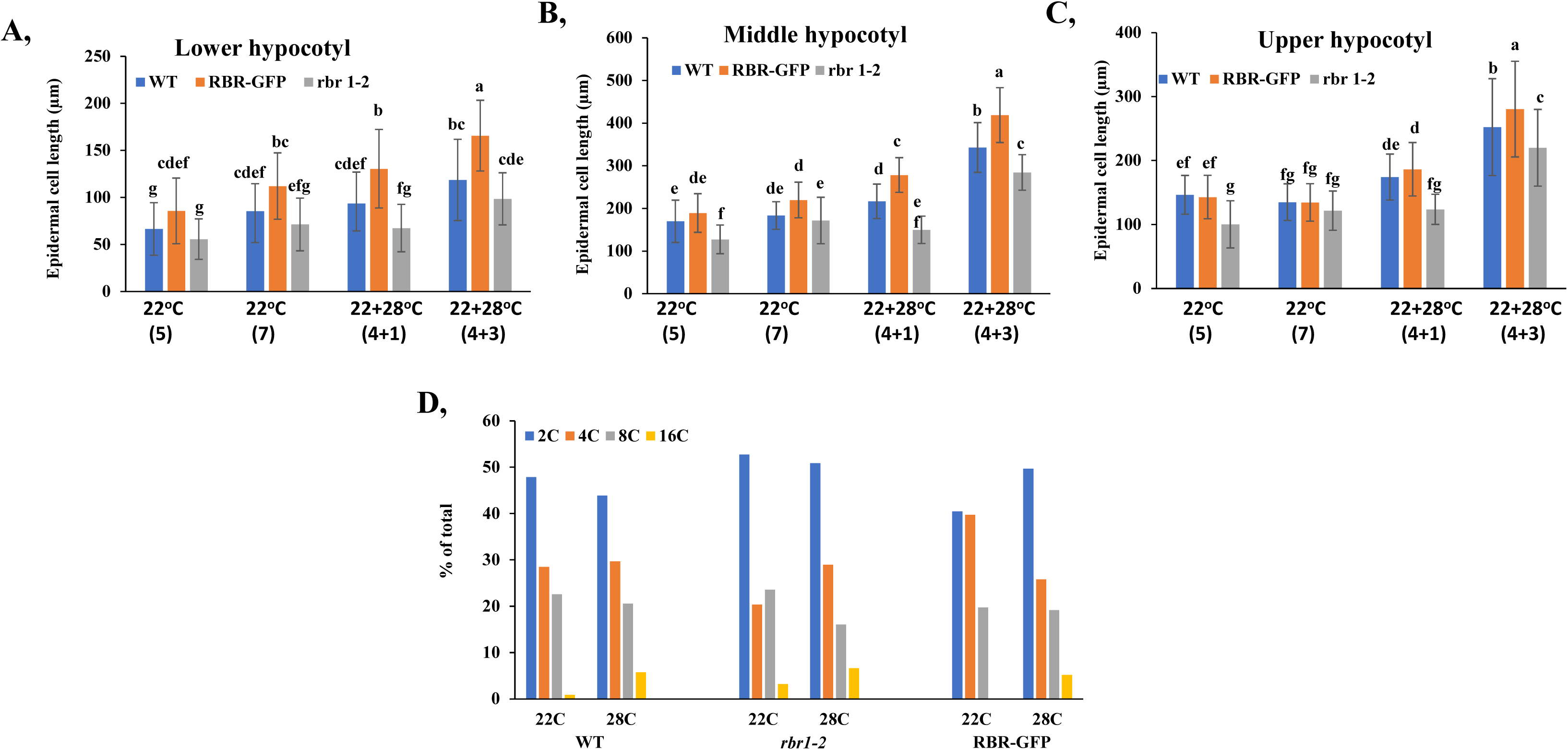
Ectopic RBR level increases epidermal size of the hypocotyl with warm temperature without affecting the ploidy level. Wilde type (WT), ectopic RBR-GFP expressing, and *rbr1-2* mutant seedlings were grown continuously at 22°C (control) or transferred from 22°C to 28°C (22+28°C) for the number of days indicated in the parentheses. **A-C**) Epidermal cells of the hypocotyls were measured in three regions (upper-A, middle-B and the bottom-C) for seedlings grown at 22°C or after transfer to 28°C as indicated. Different letters mean statistically significant differences (p < 0.05) based on one-way ANOVA analyses with Tukey’s HSD post-hoc test. **D**) The ploidy level of the hypocotyls was determined by a flow cytometer for seedlings grown continuously at 22°C or 28°C for four days, respectively.

**Supplemental Figure S5.**
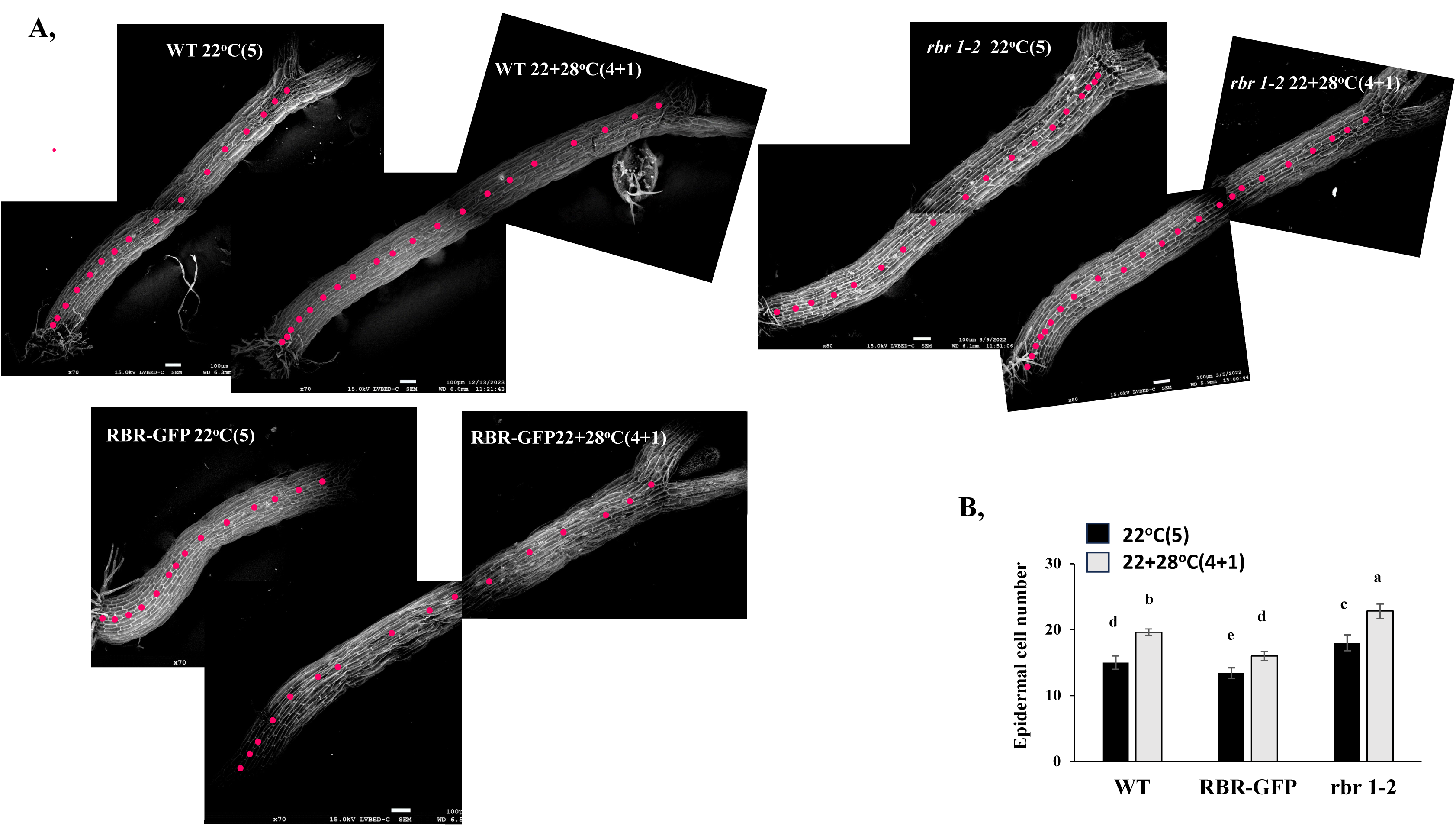
Warm temperatures increase cell numbers in the hypocotyl. Wild type (WT), ectopic RBR-GFP expressing, and *rbr1-2* mutant seedlings were grown continuously at 22°C (control) or transferred from 22°C to 28°C (22+28°C) for the number of days indicated in the parentheses. **A**) SEM images of hypocotyls of five-day-old WT, *rbr1-2*, and RBR-GFP seedlings grown at a continuous 22°C or transferred to 28°C at four days. Pink dots mark epidermal cells in the same longitudinal file. B) The number of epidermal cells in the hypocotyls developed under the same conditions as in A were counted and plotted (n=8). Different letters mean statistically significant differences (p < 0.05) based on one-way ANOVA analyses with Tukey’s HSD post-hoc test.

**Supplemental Figure S6.**
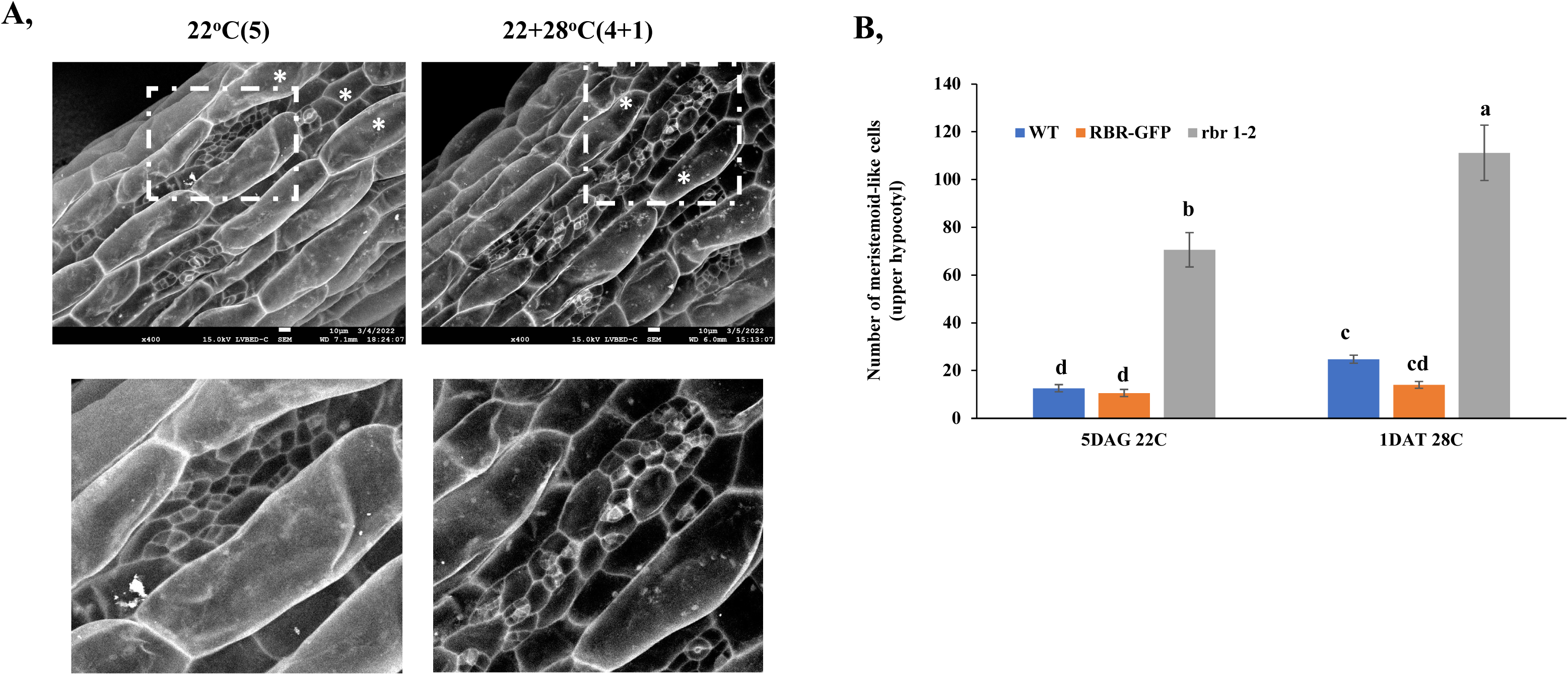
Stomatal meristemoid cells in the *rbr1-2* mutant hypocotyl show increased proliferation activity under warm temperatures, resulting in the formation of small cell clusters. **A**) Representative SEM picture of the hypocotyl of five-day-old *rbr1-2* seedlings grown at continuous 22°C (22°C(5)) or transfered to 28°C on the fourth day (22+28°C(4+1)). Meristemoid cell files can be observed between the protruding files of elongated epidermal cells indicated by asterisks. B) The number of stomatal meristemoid-like cells is highest in the *rbr1-2* mutant line compared to the wild-type (WT) and the RBR-GFP expressing lines. Warm temperature further increases the number of meristemoid cells in this line (n=10). Different letters mean statistically significant differences (p < 0.05) based on one-way ANOVA analyses with Tukey’s HSD post-hoc test.

**Supplemental Figure S7.**
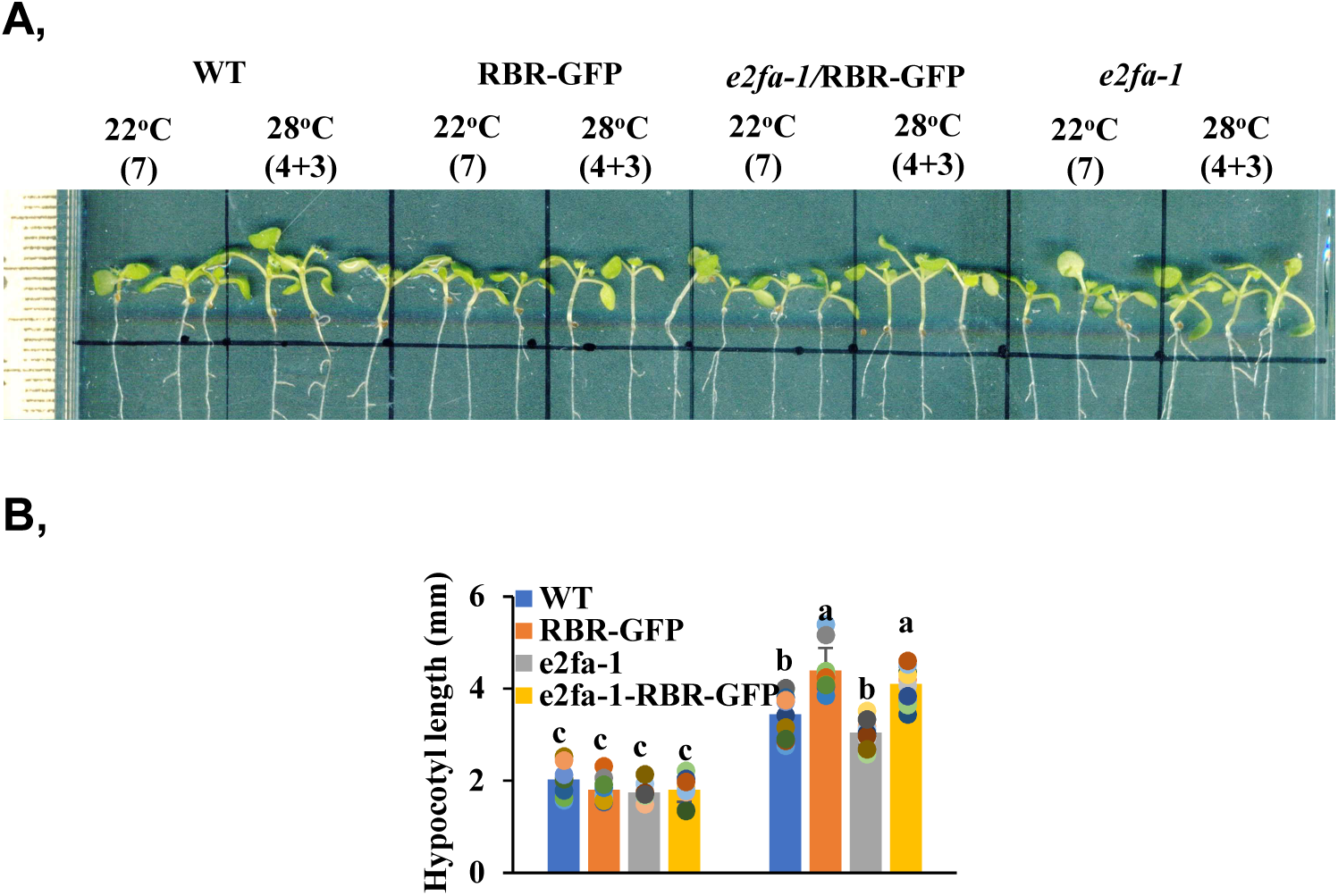
Hypocotyl length of the RBR-GFP line was not influenced by the *e2fa* mutation. **A**) Representative pictures of seedlings of the indicated genotypes grown at different temperatures for the indicated periods. **B**) Hypocotyl length was measured and plotted for the seedlings shown on A. N=5. Different letters mean statistically significant differences (p < 0.05) based on one-way ANOVA analyses with Tukey’s HSD post-hoc test.

**Supplemental Figure S8.**
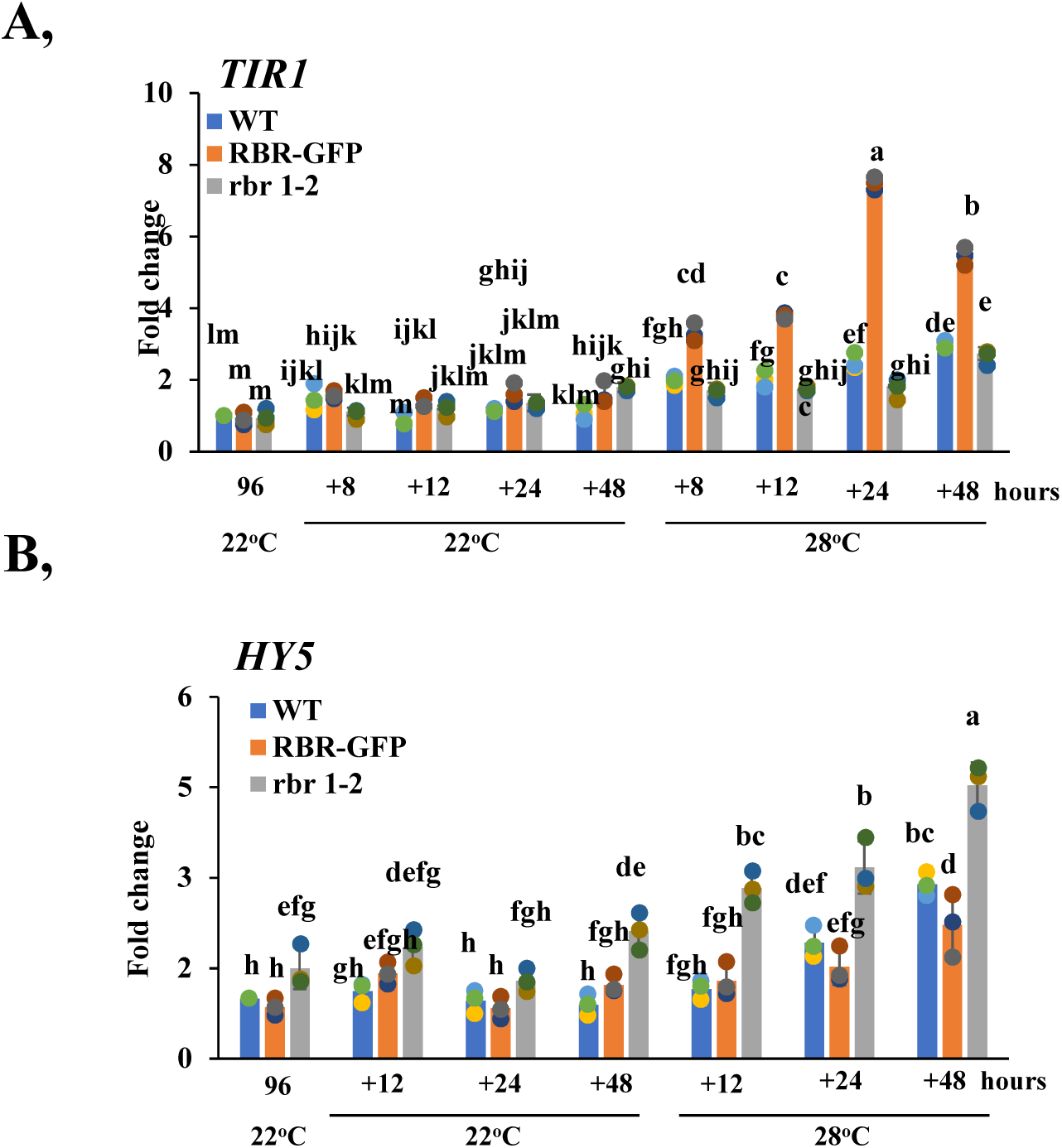
RBR oppositely regulates auxin receptor gene *TIR1* and the photomorphogen *HY5* at warm temperatures. **A-B**) Expression of *TIR1* and *HY5* was monitored by qRT-PCR at different time points after transfer of seedlings grown at 22°C for 96 hours to 22°C or 28°C as indicated. **A**) The auxin receptor F-box protein TIR1 was induced by warm temperatures, and its expression was further stimulated by ectopic RBR. **B**) Expression of *HY5* was also increased by warm temperatures but the reduced RBR level in the *rbr1-2* mutant resulted in a higher transcript level, while the ectopic RBR in the RBR-GFP seedlings suppressed the effect of warmth. Values represent fold change normalized to the value of the relevant transcript of the seedlings at T0 (96 hours), which was set arbitrarily at 1. Data are means +/- sd. N=3 biological replicates. Different letters mean statistically significant differences (p < 0.05) based on one-way ANOVA analyses with Tukey’s HSD post-hoc test.

**Supplemental Table S1.**
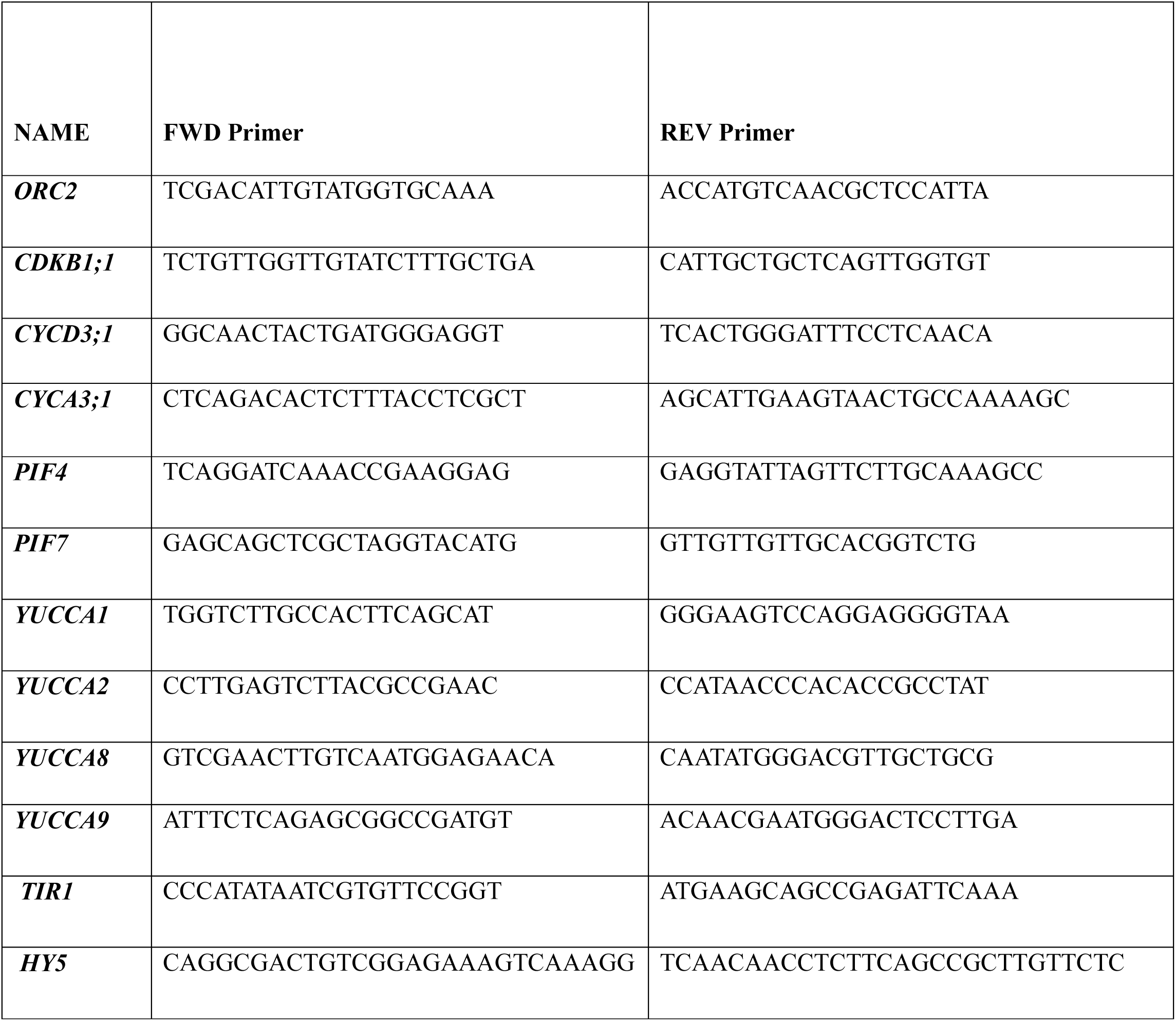
List of primers and their sequences used for qRT-PCR analysis. In the first row from left to right, gene identification (GI), gene’s name, forward (FWD) and reverse (REV) sequences.

